# The DeepTune framework for modeling and characterizing neurons in visual cortex area V4

**DOI:** 10.1101/465534

**Authors:** Reza Abbasi-Asl, Yuansi Chen, Adam Bloniarz, Michael Oliver, Ben D.B. Willmore, Jack L. Gallant, Bin Yu

**Affiliations:** Department of Electrical Engineering and Computer Sciences; Department of Statistics; Helen Wills Neuroscience Institute; Department of Psychology, University of California, Berkeley, CA 94720

## Abstract

Deep neural network models have recently been shown to be effective in predicting single neuron responses in primate visual cortex areas V4. Despite their high predictive accuracy, these models are generally difficult to interpret. This limits their applicability in characterizing V4 neuron function. Here, we propose the DeepTune framework as a way to elicit interpretations of deep neural network-based models of single neurons in area V4. V4 is a midtier visual cortical area in the ventral visual pathway. Its functional role is not yet well understood. Using a dataset of recordings of 71 V4 neurons stimulated with thousands of static natural images, we build an ensemble of 18 neural network-based models per neuron that accurately predict its response given a stimulus image. To interpret and visualize these models, we use a stability criterion to form optimal stimuli (DeepTune images) by pooling the 18 models together. These DeepTune images not only confirm previous findings on the presence of diverse shape and texture tuning in area V4, but also provide rich, concrete and naturalistic characterization of receptive fields of individual V4 neurons. The population analysis of DeepTune images for 71 neurons reveals how different types of curvature tuning are distributed in V4. In addition, it also suggests strong suppressive tuning for nearly half of the V4 neurons. Though we focus exclusively on the area V4, the DeepTune framework could be applied more generally to enhance the understanding of other visual cortex areas.

## 1 Introduction

Understanding the function of primate visual pathways is a major challenge in computational neuroscience. Along the ventral visual pathway, cortical area V4 is of particular interest. It is a large retinotopically-organized area located intermediate between the early primate visual cortex areas such as V1 and V2 and high-level areas in the inferior temporal (IT) lobe. V4 is believed to be crucial for visual object recognition and visual attention, but its functional role remains mysterious. Computational studies of primary visual cortex have produced powerful quantitative models of V1 [5]. Contrastingly, area V4 is more difficult to model computationally than V1. This is mainly due to its highly nonlinear response [42] and diverse tuning properties [30].

To understand the tuning properties of V4 neurons, one dominant traditional approach is to use handcrafted and limited synthetic stimuli (e.g. [10, 28]) to probe V4 neurons. For example, by comparing V4 neuron responses to Cartesian gratings with those to polar and hyperbolic (non-Cartesian) gratings, Gallant et al. [10, 11] found that V4 neurons are most selective for non-Cartesian gratings containing multiple orientations. Through a parameterized set of contour stimuli varying in angularity, curvature, and orientation, Pasupathy and Connor [27, 28] discovered that V4 neurons are selective to curved contour features. While such studies have successfully quantified V4 neuron responses to synthetic shapes, the tuning properties of most V4 neurons cannot be fully explored through these limited sets of stimuli [30].

An alternative approach to designing synthetic stimuli is using a large collection of natural images directly as stimuli. This approach reduces the difficulty in stimuli design, but creates a huge challenge in modeling. Specifically, it has been found that previously proposed simple and shallow computational models of V4 neurons perform poorly on natural images [8, 25, 30]. For instance, David et al. [8] introduced the spectral receptive field (SRF) model to account for second order nonlinear response properties. The SRF model enhances our understanding of V4 orientation tuning properties, but its average prediction performance for the V4 neurons studied is far from satisfying [30]. More recently, advances in deep convolutional neural networks (CNNs) with multiple layers of linear and non-linear operations have led to more accurate predictive models for neurons in V4 and IT [48, 4, 47]. While this deep, convolutional and non-linear architecture is the key to the high predictive performance, it also makes the models difficult to interpret. This limits their usefulness in advancing neuroscience. A natural question arises: can we use these complex and accurate models to infer tuning properties of V4 neurons?

In this paper, we propose the DeepTune framework as a tool to visualize and interpret predictive models of single neurons. In order to make the interpretations be less dependent on arbitrary neural network architecture choices, we build an ensemble of 18 CNN-based models per neuron instead of a single model. The models vary in architecture, but all have comparable high and state-of-the art prediction accuracies. Each model uses a CNN to extract features from an input image. The CNN is pre-trained to perform object classification on the ImageNet dataset [32]. The extracted features are then used as predictors to train a regularized linear regression model with the neuron firing rate as the response. This approach of applying a pre-trained model to a new prediction task is known as transfer learning [36]. For each neuron, we then generate DeepTune images that are obtained via gradient optimization of the fitted models. Aggregating the DeepTune generation process from 18 models via a stability criterion, we further introduce the consensus DeepTune images for each neuron. We show that interpreting the components of DeepTune images that are consistent across 18 models and the consensus one can help better characterize the tuning property of a neuron and gain robustness against modeling choices. Finally, we perform population analysis of all DeepTune images from 71 neurons to illustrate the curvature tuning diversity and suppressive tuning in V4.

### Significance Statement

Understanding how primates process visual information and recognize objects in an image is a major problem in neuroscience. Along the visual pathway, the midtier cortical area V4 is of particular interest. Despite its importance in the hierarchical organization of visual processing, its function remains elusive. Accurate deep neural network-based predictive models are built for responses of V4 neurons to natural image stimuli. While interpreting these models is traditionally difficult, we introduce the DeepTune framework to equip these complex models with stable interpretation and visualization. The DeepTune images provide rich, concrete and naturalistic characterizations of V4 neurons that refine significantly findings of previous studies. They hold promise as better natural input stimuli for future closed-loop experiments.

## 2 CNN-based models are highly predictive of V4 neuron responses on natural stimuli

We have recorded firing rates of 71 well isolated neurons in V4 from two awake-behaving male macaques. These recordings were previously used to study the sparseness of neural codes in the area V4 (but without predictive models) [44]. The stimuli consist of a random sample of circular patches of grayscale digital photographs from a commercial digital library (Corel). Uniformly random sampled images without replacement were then concatenated into long sequences so that each 16.7 ms frame contained a random image from the library. When presented to the macaques, all images were centered on the estimated classical receptive field (CRF, see *SI Data Collection* for CRF estimation procedure). The image size was adjusted to be two to four times the CRF diameter (Figure 1-C). The training data set for each neuron contains 8,000-24,000 natural images (4,000-12,000 distinct ones * 2). Spike counts were measured at 60Hz, resulting in two measurements per image. For the holdout test dataset, 600 images (300 distinct ones * 2) were shown for each neuron in a fixed order, distinct from the images shown for the training dataset. The sequence of test images was repeated; for each neuron, each image in the test dataset was shown 8-10 times. The resulting spike counts were averaged to provide a more precise estimate of the expected spike count. In addition, repeats also allowed for estimating the amount of variance in the neuron explainable by the stimulus image [35] (see *SI Data Collection* for details).

**Figure 1:**
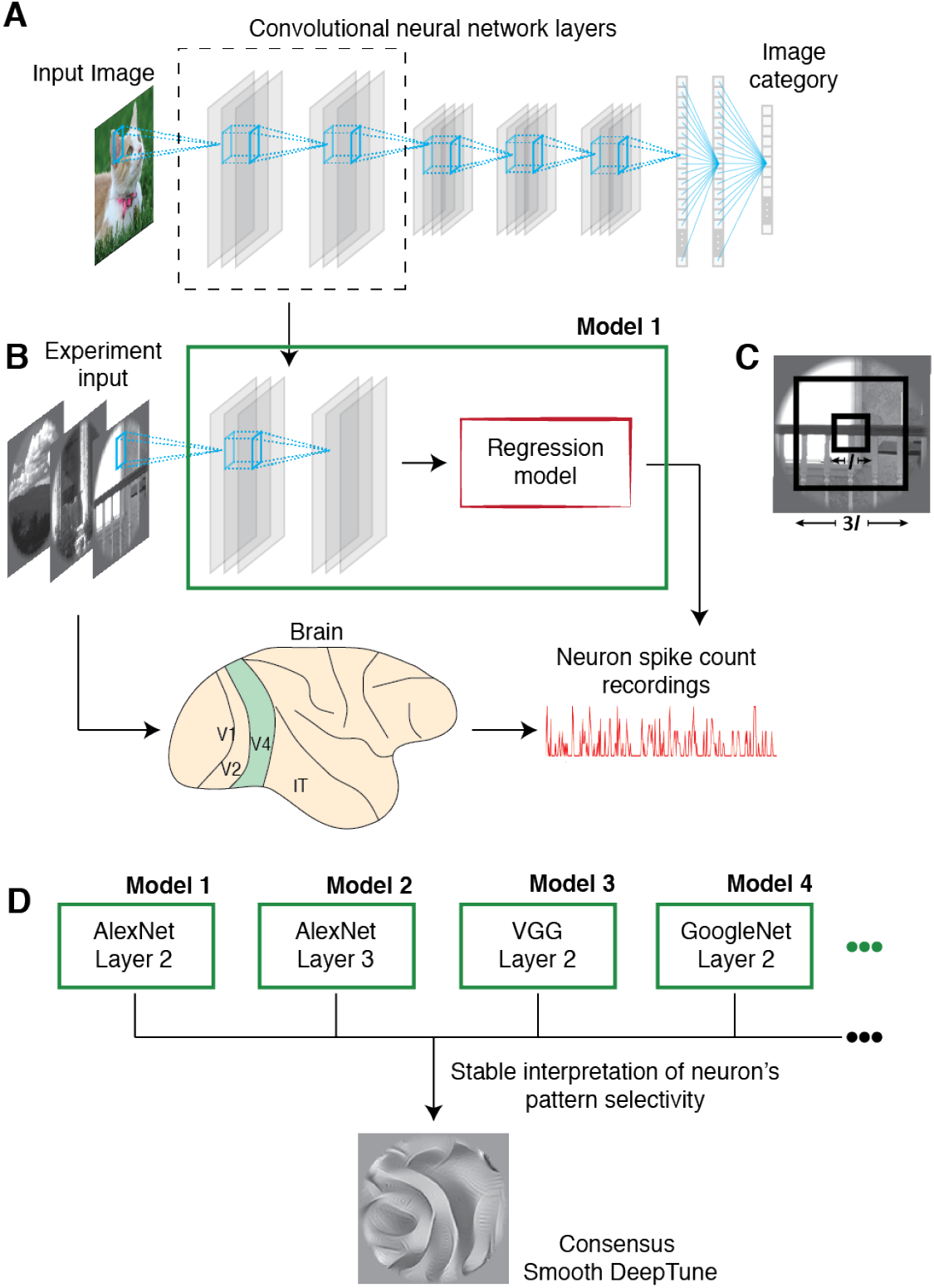
DeepTune framework through transfer learning: first, we use features from pre-trained convolutional neural networks (CNNs) in regularized regression to predict (spike) firing rates of neurons in the visual area V4; second, stability-driven DeepTune images across 18 CNN-based predictive models are generated for interpretation. **A.** Architecture of a convolutional neural network (CNN) pre-trained to perform 1000-class image classification task on the ImageNet dataset (e.g. AlexNet). **B.** An input image is propagated forward in a fixed layer of the CNN, yielding a feature vector representation of the image. This vector is used to fit a regularized linear regression model to predict firing rates of each V4 neuron. **C.** The classical receptive field (CRF) during the experiment is set in the middle of the stimuli with width *l* while the whole image has the width 3*l*. **D.** 18 accurate predictive models are obtained using features from layers 2, 3, 4 of three pre-trained AlexNet, GoogleNet, VGG, with either *ℓ*_1_ (lasso) or *ℓ*_2_ (ridge) regularized linear regression. DeepTune, a stability-driven interpretation and visualization framework of CNN-based model (across multiple such models) is proposed to characterize V4 neurons’ tuning preferences (more details in the Results section 2). The consensus DeepTune image for one neuron (corresponds to Neuron 1 in Figure 3-A) is shown and displays a stable curvature pattern with edges forming an approximately ninety-degree angle.

We introduce a transfer learning framework (Figure 1) to build predictive models in two stages for our V4 stimulus-response data as just described. For a given layer of a pre-trained CNN and for each input stimuli, in the first stage (Figure 1-A), we extract intermediate outputs from that layer of CNN as features. In the second stage (Figure 1-B), these features serve as predictors in a regularized linear regression (such as Ridge [13] or LASSO [41]) with time-lagged spike rates as the responses. Specifically, for one stimulus image at time *t* denoted as **z**_*t*_ ∈ ℝ^*s*×*s*^ (*s* = 227 in the AlexNet CNN model [19]), the given layer of CNN transforms this image into a flattened feature vector **x**_*t*_ ∈ ℝ^*d*^ (*d* = 256 *×* 13 *×* 13 in the AlexNet-Layer2 CNN model). This feature transform is denoted as function *h* : ℝ^*s*×*s*^ ↦ ℝ^*d*^. Since the responses of V4 neurons to a sequence of images are sensitive to the recent history of images shown to the subject, we build the models with multiple time lags. More precisely, we regress *y_t_* against the training image features from last *k* frames of video prior to and including time *t*, i.e. **z**_*t*_, …, **z**_*t−k*+1_. The time lag *k* is set to be 9 (consisting frames at 0, 16.7, …, 133.6 ms) as in previous studies with similar data recordings (e.g. [8, 45]). Finally, our predictive model for a single neuron response takes the following form

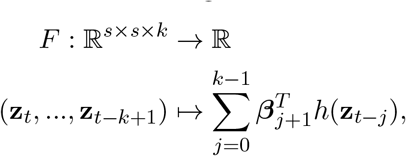

where (*β*_1_, …, *β_k_*) ∈ ℝ^*d*×*k*^ are the regression parameters to be estimated and *h* is the fixed CNN feature transform. The model parameters are learned by solving the following regularized linear regression problem

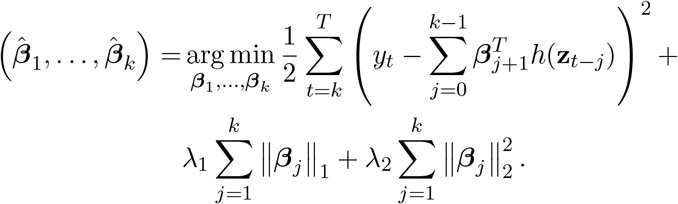

If not specified in the rest of the paper, the regularization is taken to be *ℓ*_2_ norm by setting λ_1_ = 0 (Ridge). The analysis with *ℓ*_1_ norm regularization (LASSO) by setting λ_2_ = 0 to enforce sparsity is discussed in *SI Stability of Analysis*.

The CNNs used are pre-trained CNNs for classification tasks. They are trained based on a 1000-object classification task on the ImageNet dataset from the ImageNet Large Scale Visual Recognition Challenge [32]. One legitimate concern of deploying neural networks in modeling is that interpretations about the models may depend on the details of the neural network architecture choices. To address this problem, we use three different neural network architectures to model V4 neurons: AlexNet [19], GoogleNet [39] and VGG [38]. All three networks have high classification performance on ImageNet recognition challenge and are known to provide transferable image features in other computer vision tasks such detection and segmentation [36, 49]. To vary the number of layers, we use features from layer two, three and four of each network. Later in this section, we show that using layer 1 and layers higher than layer 4 leads to lower prediction accuracies or has too large receptive fields not comparable with those of V4 neurons. Finally, on top of the CNN features, either Ridge or Lasso regression is used to predict the (spike) firing rates. As a result, we obtain 18 models for each neuron (3 nets *×* 3 layers *×* 2 regression models). Next we provide detailed prediction performance of these 18 models and compare them to previous models in the literature before we propose the stability-driven interpretation and visualization framework of DeepTune based on a stable aggregation of all 18 models.

To determine quantitatively how well our models describe the responses of each neuron, we test their performance on the holdout test set. All our models were estimated using the training data set. The correlation between the firing rates predicted by the model and the actual average firing rates on the test set is used as the prediction performance for all our 18 models. As a baseline for comparison, we also fit a V1-like Gabor wavelet model [7, 16]. The Gabor wavelet model first extracts image features by applying a bank of linear Gabor wavelet filters to the input image at varying orientations, spatial frequencies and phases, followed by half-wave rectification and a compressive nonlinearity, then regresses the responses of each neuron using Ridge regression [13].

Our AlexNet-Layer2 (+Ridge) model has a average correlation coefficient of 0.44 (or 0.52 for noise-corrected correlation coefficient [35]) on the holdout test set. It achieves the state-ofthe-art prediction accuracy for V4 neurons on natural image stimuli [8, 46]. Comparing to [8], our average correlation coefficient is about 0.15 higher. As shown in Figure 2-D, all of the 18 models have average correlation coefficients higher than 0.42. For nearly all of the 71 V4 neurons, they are all more accurate than the V1-like Gabor wavelet model (with an average correlation coefficient 0.33). Due to space limitations, we plot the results only for 4 models, which are all based on AlexNet-Layer2, AlexNet-Layer3, VGG-Layer2, GoogleNet-Layer2 (and ridge) in Figure 2-A and 2-B. The first two models are chosen in order to demonstrate stability of prediction results and interpretations across different CNN layers, while the other two models are chosen to show stability across different CNN architectures. In Figure 2-C, we compare the average prediction performance for models from all 7 layers of AlexNet for 71 neurons. The model based on AlexNet-Layer1 has similar performance to that of the V1-like Gabor wavelet model; while models from layers 2 to 5 have much higher predictive performance (e.g. 0.44 for layer 2, 0.46 for layer 5). This justifies the recent finding [36] that the intermediate layers of pre-trained CNNs (on large-scale image classification tasks), like AlexNet, can extract more complex features than the first layer and Gabor wavelets.

**Figure 2:**
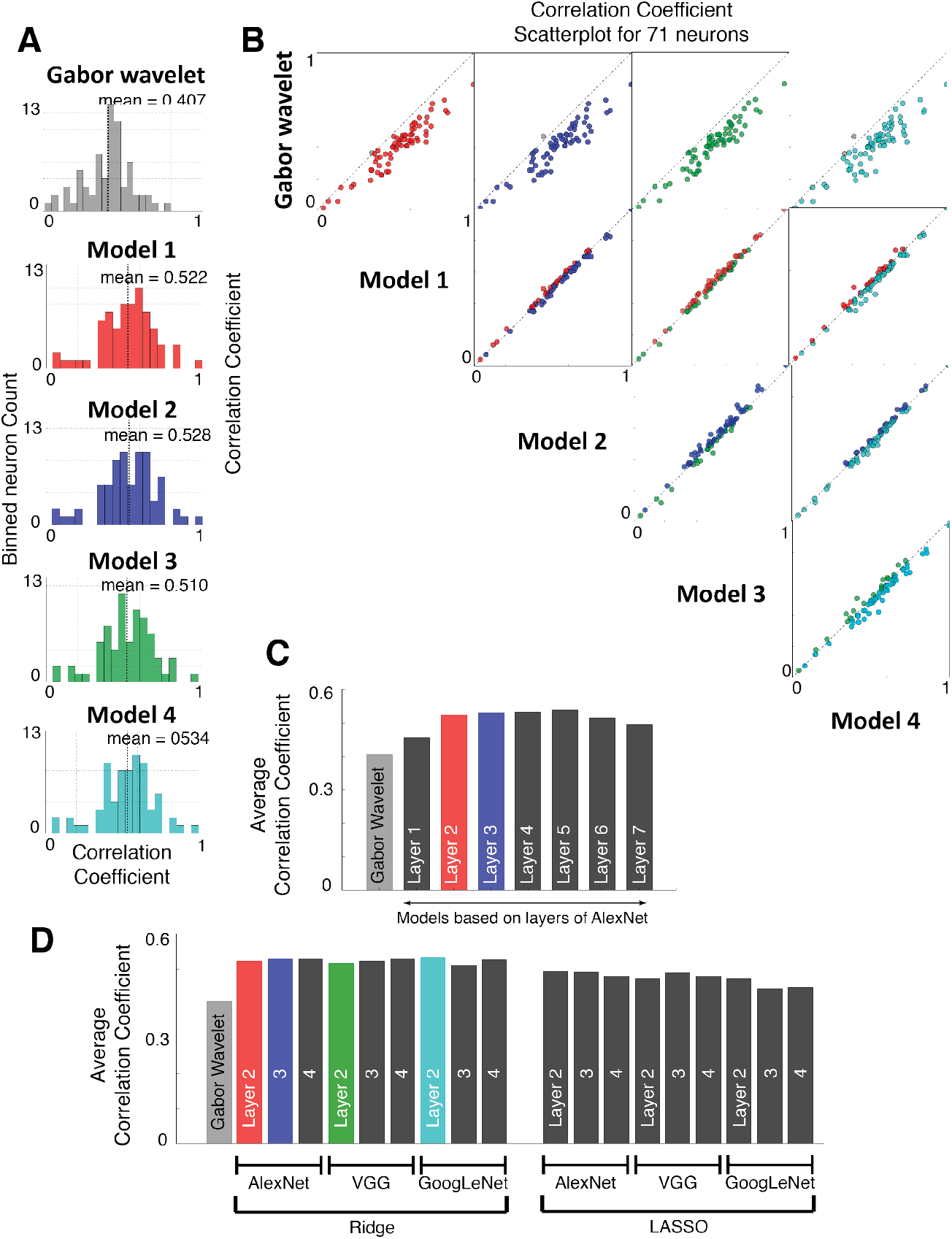
CNN-based models outperform V1-like Gabor wavelet model in terms of noise-corrected correlation coefficient [35] as the prediction performance measure. **A.** Histogram of noise-corrected correlation coefficients over population of 71 V4 neurons for five models are shown, where baseline model is V1-like Gabor wavelet model, Model 1 is AlexNet-Layer2, Model 2 AlexNet-Layer3, Model 3 VGG-Layer2, and Model 4 GoogleNet-Layer2. Ridge regression is used in all 4 models. **B.** Scatter plots comparing each pair among Models 1-4. Results for the other 14 models are shown in *SI Stability of Analysis, Figure S6*. **C.** The average prediction performance across 71 neurons for models from all 7 layers of AlexNet with ridge regression. The model based on AlexNet-Layer1 has similar performance to that of the V1-like Gabor wavelet model; while models from layers 2 to 5 have higher predictive performance. **D.** The average prediction performance across 71 neurons for all 18 models. All 18 models perform similar in prediction and much better than the Gabor wavelet model.

In order to be consistent with the literature [29, 46, 8], we also report the proportion of explainable variance captured by a model. It attempts to control for differences in noise levels between experimental setups, individual neurons, and brain regions. We estimate the explainable variance through the noise-corrected correlation coefficient [35] using the repeated data in the holdout set (see *SI Methods* for more information). Averaged over the 71 V4 neurons, the AlexNet-Layer2 and ridge model captures 30.3% of the explainable variance. This performance matches the 30% of computational models for area V2 [45]. The unexplained portion of the response is very likely to have resulted from two factors: visual tuning properties not described by the AlexNet-Layer2 (and ridge) model and non-stimulus influences on the response. The latter is unlikely to be removed completely given our experimental setups [45]. Note that the prediction task on the natural images in this paper is substantially harder than that on images with artificial objects overlaid in [48]. Besides this work [48] on simpler natural image stimuli, our CNN-based models demonstrate a large improvement in prediction performance over previous works with natural image stimuli similar to ours [8, 46]. In the next section, we take advantage of this high prediction accuracy to better characterize of V4 tuning properties via DeepTune images.

## 3 DeepTune as a naturalistic visual representation of tuning

It has long been challenging to fully characterize shape tuning properties in area V4. There are two main difficulties: the absence of highly predictive and biologically plausible computational models for the nonlinear response properties of V4 [30], and the lack of systematic methods to generate relevant complex natural stimuli to probe V4 neurons more efficiently. Given the state-of-the-art predictive performance of our CNN-based models, it is natural to ask whether these models could also provide a better characterization of shape tuning (e.g. angular, curvature or orientation tuning) or texture tuning in area V4. However, unlike existing studies using relatively simple Gabor wavelets [7, 16] or Fourier transform [8], complex nonlinear CNN features in our models make it extremely challenging to consistently interpret our models.

Inspired by computer vision advances in visualizing CNNs [51, 21], we introduce *DeepTune images* as a naturalistic visual representation of tuning for a V4 neuron. The DeepTune images are made of a collection of reconstructed images that jointly represent the shape tuning properties of a neuron. For each neuron and for each given model, a *DeepTune image* (or preferred DeepTune image) is obtained by optimizing over the input image space to maximize a regularized model output (predicted neuron response). Starting from a random image (e.g. white noise image with zero mean and fixed small variance), we use the gradient ascent method to gradually increase the model output until convergence. Formally, given a fixed predictive model at a particular time lag (the single lag time that causes best prediction performance in a 10% validation set split of the training set) *f* : ℝ^*s*×*s*^ ↦ ℝ, we seek an input image **z** ∈ ℝ^*s*×*s*^ that minimizes the following objective function:

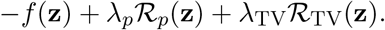

The regularization terms are included to capture prior information about natural images. That is, the optimization search is constrained to be close to the set of smooth and naturalistic images [21]. The specific regularization choices above are motivated by image denoising techniques [31] and by natural image statistics [37]. The first regularizer *ℛ_p_* (the *ℓ_p_*-norm of a vectorized image pixels) encourages the intensity of pixels to stay small. By choosing a large *p* (*p* = 6 in our analysis), this regularizer prevents the solution image from taking extremely large pixel values. The second regularizer *ℛ*_TV_ controls the total variation norm of an image. It encourages the image to be smooth and removes excessive high-frequency details (see *SI Methods* for more information).

The collection of DeepTune images is constructed from all 18 predictive models. In addition, we verify that 10 independent random initializations of starting images do not change the output much (see *SI Stability of Analysis, Figure S7*). Similarly, an inhibitory DeepTune is obtained by minimizing instead of maximizing the model output. We note that the DeepTune images differ from the traditional receptive fields in neurophysiology [14, 7] in two ways: multiple images are used to describe tuning properties of a single neuron; they are more naturalistic representations of tuning with a higher resolution.

Figure 3-A shows the DeepTune images from 4 of our 18 models built for Neuron 1. We visually observe that these DeepTune images share a stable curvature pattern with edges forming an angle of nearly 90 degrees. The rest 14 DeepTune images produced from the other 14 models differ slightly, but the main curvature pattern remains relatively stable (see *SI Stability of Analysis Figure S8*). That is, the curvature angle stays close to 90 degrees and the spatial location of the curvature pattern remains at left side of the image. To further quantify the curvature angle and spatial frequency, we compare the power spectral densities (PSD) of these DeepTune images in Figure 3-B. All four DeepTune images share a strong and stable frequency component in the range of 45 to 135 degrees with spatial frequencies of 2 to 5 cycles per receptive field (green). Note that the high frequency components from the Model-4 DeepTune image are not consistent with the other three models. Especially, GoogleNet-Layer2 model has high frequency components that are not present in three other models. Therefore these components likely reflect noise and should be discounted. In Figure 3-C, we visualize the spectral receptive field (SRF) model [8] for Neuron 1. The SRF visualization shows the frequency components of the stimulus image selected by SRF model. The color map (red-blue) is chosen to be different from that of the DeepTune Fourier transform (green-pink). The color map difference serves a reminder of the difference between PSD and SRF. As observed from the DeepTune image PSD, the SRF model also shows that Neuron 1 exhibits a strong preference to the frequency component in the range of 45 to 135 degrees with spatial frequency of 2 to 5 cycles per receptive field. In addition to DeepTune and SRF, this curvature tuning is further supported by the curvature patterns in the images from training and test sets with the highest responses for Neuron 1 (Figure 3-D and E). Figure 3-E illustrates the measured and predicted firing rates in test set from the 4 models as well as the predicted firing rates from the SRF model. For this Neuron 1, our 4 models have similar prediction accuracies (correlations on the holdout set between 0.61 to 0.64), while the SRF model has difficulty capturing the peak firing rates as seen in the lower plot of Figure 3-E, with a corresponding correlation of 0.42.

**Figure 3:**
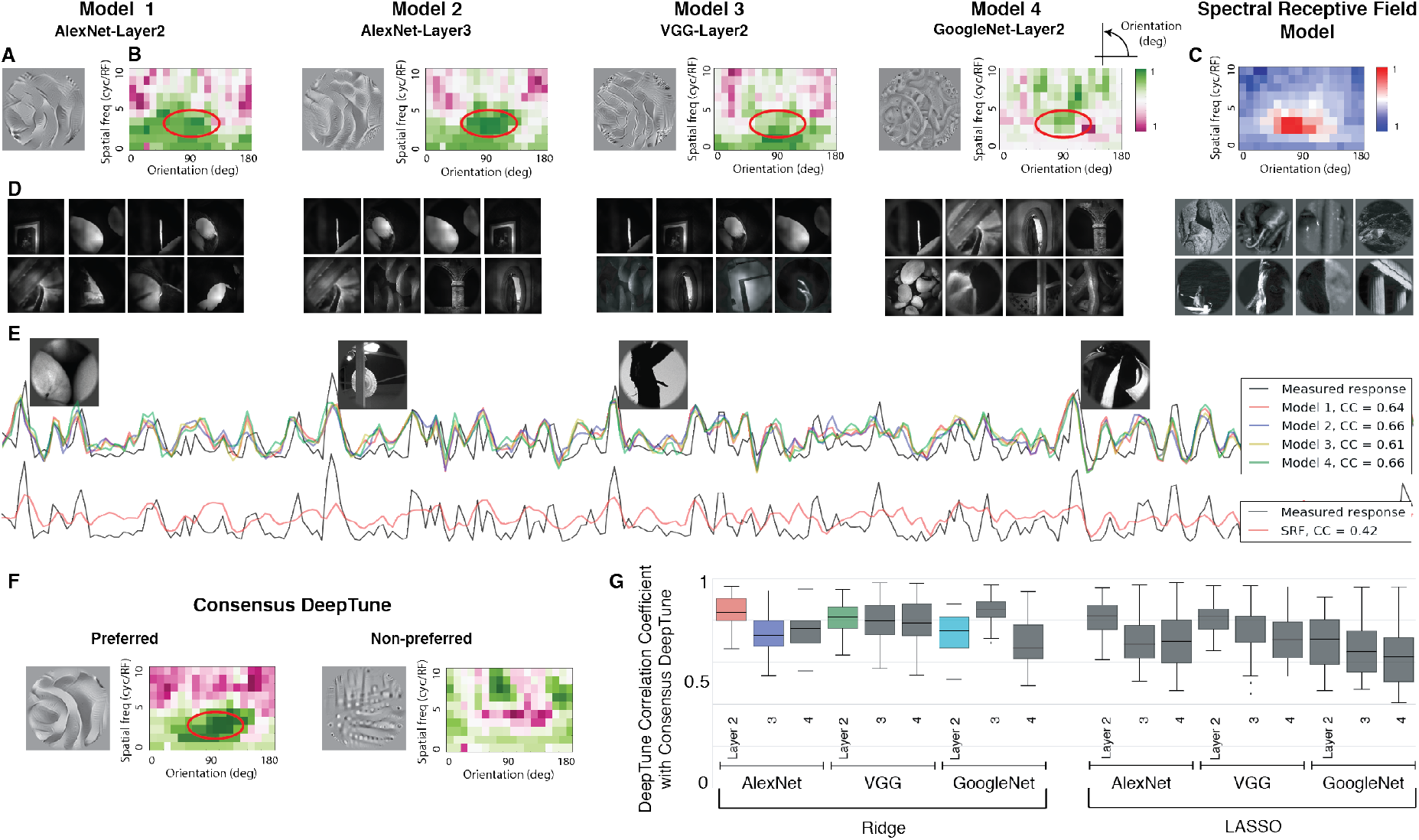
DeepTune images from four of our 18 models built for Neuron 1. **A.** DeepTune images based on Models 1-4 for Neuron 1. These images share a visually stable curvature pattern with edges forming an approximately ninety-degree angle. **B.** Power spectral densities (PSDs) of the DeepTune images in polar coordinates. Through the PSDs, all four DeepTune images share a strong and stable frequency component in the range of 45 to 135 degrees with spatial frequency of 2 to 5 cycles per receptive field (the green color). **C.** Visualization of spectral receptive field (SRF) [8] model for Neuron 1. The SRF visualization emphasizes in red the frequency components of the stimulus image selected by the SRF model. The pattern selectivity according to SRF is consistent with the stable part of the PSDs of DeepTune images (highlighted in red circles). **D.** Images from training set with the highest responses for Neuron 1. Similar curvature patterns to the DeepTune visualization are visible in these images. **E.** The measured and predicted (spike) firing rates in the test set from Models 1-4 as well as the SRF model for Neuron 1. Images from the test set with the highest responses are visualized on top of the corresponding spike rate. Similar curvature patterns are visible in these images. Correlation coefficients between the measured and predicted firing rates are shown in the right panel. All four models outperform the SRF model. **F.** The consensus DeepTune image for Neuron 1. Both excitatory, inhibitory DeepTune images and the corresponding PSDs are shown. The excitatory pattern based on the consensus DeepTune exhibits the curvature contour that is similar to those from the four models in panel A. The inhibitory pattern visually consists of lines orthogonal to the preferred curvature contour, confirmed via PSD visualization on the right. **G.** Each box-plot corresponds to a CNN-based model among the 18 models and is based on 71 raw-pixel correlation coefficients. Each such coefficient corresponds to a neuron and is calculated between the consensus DeepTune image and a DeepTune image from that model and for that neuron. DeepTune images from AlexNet-Layer2 and GoogleNet-Layer 3 have the highest similarity on average to the consensus DeepTune image

In addition to the visual comparison of 18 distinct DeepTune images generated from 18 models, we introduce consensus DeepTune to capture in a single image the stable patterns across 18 models. The consensus DeepTune image is obtained via a similar optimization scheme as in the original DeepTune optimization for a single model, but with an aggregation of gradient information from all 18 models. The aggregated gradient maintains the stable components in the gradients and discounts the unstable components (more details in *SI Methods*). Both excitatory and inhibitory consensus DeepTune images for Neuron 1 are shown in Figure 3-F. The excitatory consensus DeepTune (Figure 3-F) exhibits curvature contour patterns that visually matches all 4 models (Figure 3-A). The power spectral density (PSD) to the right of the consensus DeepTune image in Figure 3-F similarly matches the individual models. This PSD displays strong frequency components in the range of 45 to 135 degrees with spatial frequencies of 2 to 5 cycles per receptive field. On the other hand, the inhibitory consensus DeepTune consists of lines orthogonal to the curvature contour (see *SI Stability of Analysis* for comparison with inhibitory DeepTune images from all 18 models). Some blobs are also visible in the inhibitory consensus DeepTune image, suggesting that the response of Neuron 1 is attenuated by blob-like texture patterns. This is further supported by observing that the inhibitory PSD contains strong high frequency components on the top center.

The consensus DeepTune image captures the stable components of DeepTune images across our 18 models. It can be visually observed that the DeepTune images from a number of individual models are very similar to the Consensus DeepTune (see *SI Stability of Analysis, Figure S8*). To quantify this similarity, we compute the Pearson correlation coefficient between pixel values of the consensus DeepTune and those of each DeepTune image. Figure 3-G visualizes boxplots of these correlation coefficients. Each boxplot corresponds to one of the 18 models and shows the distribution of 71 correlation coefficients for all 71 neurons for this model. The median correlations for all of the models are considerably high. The highest median correlation is 0.83 which is achieved by AlexNet-Layer2 and GoogleNet-Layer3 with ridge regression. Models with lasso tend to have lower similarities to the consensus DeepTune. Due to space limitations, in the subsequent sections we present by default the consensus DeepTune image as a stable representation of a V4 neuron’s tuning property. Although a single consensus DeepTune image seems to be sufficient, the stability analysis across 18 DeepTune images are necessary to determine the spatial locations of the stable parts. This is to ensure that we identify only the stable locations of the consensus DeepTune image to be interpreted.

## 4 Model-selected CNN features highlight receptive fields

The DeepTune images described in the previous section treated the CNN-based model as an end-to-end network. In this section, we show that analyzing the intermediate stages of a CNN-based model for a neuron can provide further information. The regression weights and the CNN features are of main interest. This analysis not only provides an independent and alternative interpretation of V4 neurons, but also allows us to compare our results to previously studied spatial receptive fields of V4 neurons.

Taking AlexNet-Layer2 model as an example, we examine its regression weights (see *SI Stability of Analysis, Figure S11* for visualization of weights from other models). Regression weights with large magnitudes indicate high sensitivity of the neuron to particular image features. The AlexNet-Layer2 features are of dimension 256 *×* 13 *×* 13. They consist of 256 different convolutional filters that are spatially located on a grid of size 13 *×* 13. The corresponding regression weights at one time lag is of the same dimension. We examine the regression weights by asking the following two questions: where on the image are the regression weights with the largest magnitudes? What kinds of convolutional filters contribute the most to the prediction performance?

To answer the first question, we define an *average regression weight map* as the sum-of-squares pooling of regression weights on the CNN features. It is defined across the different convolutional filters and the time lags at each location on the 13 *×* 13 spatial grid. Formally, for each neuron, let *β*̂_*mijk*_ be the regression weight for filter *m* at spatial location (*i, j*) and lag *k*. Then the average regression weight map Φ ∈ ℝ^13*×*13^ is defined as follows:

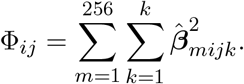

Figure 4-A shows the average regression weight map from the AlexNet-Layer2 model for 4 neurons. On the 13 *×* 13 grid map, lighter pixel color indicates higher weight map value. Maps from other models share stable shape and location (see *SI Stability of Analysis, Figure S11* for a comparison across models). For each neuron, the average regression weight map presents an estimate for the spatial receptive field. Maps for V4 neurons exhibit diverse shapes. For example, the receptive fields for Neurons 1 and 2 have round shapes, while those for neurons 3 and 4 form straight or curved band shapes. These CNN-based spatial receptive fields provide an alternative to [24] for showing diversity in the size and shape of the receptive fields of V4 neurons. These regression weight maps are also indicative of the regions where DeepTune images across 18 models share stable patterns. Figure 4-B displays the DeepTune images from the AlexNet-Layer2 model for the 4 neurons, along with the consensus DeepTune images in Figure 4-C. The corresponding inhibitory DeepTune image and consensus inhibitory DeepTune image are shown in Figure 4-D and E respectively. Looking at the patterns of the DeepTune images, Neuron 1 is tuned to the curvature-contour shapes with edges forming an approximately ninety-degree angle. Neuron 2 is tuned to blob-like patterns and textures. Neuron 3 is selective to curvature patterns with a strong diagonal line preference and Neuron 4 is tuned to corner-like shapes with edges forming ninety-degree angles. The tuning patterns shown via DeepTune are consistent with receptive field shapes shown in regression weight maps.

**Figure 4:**
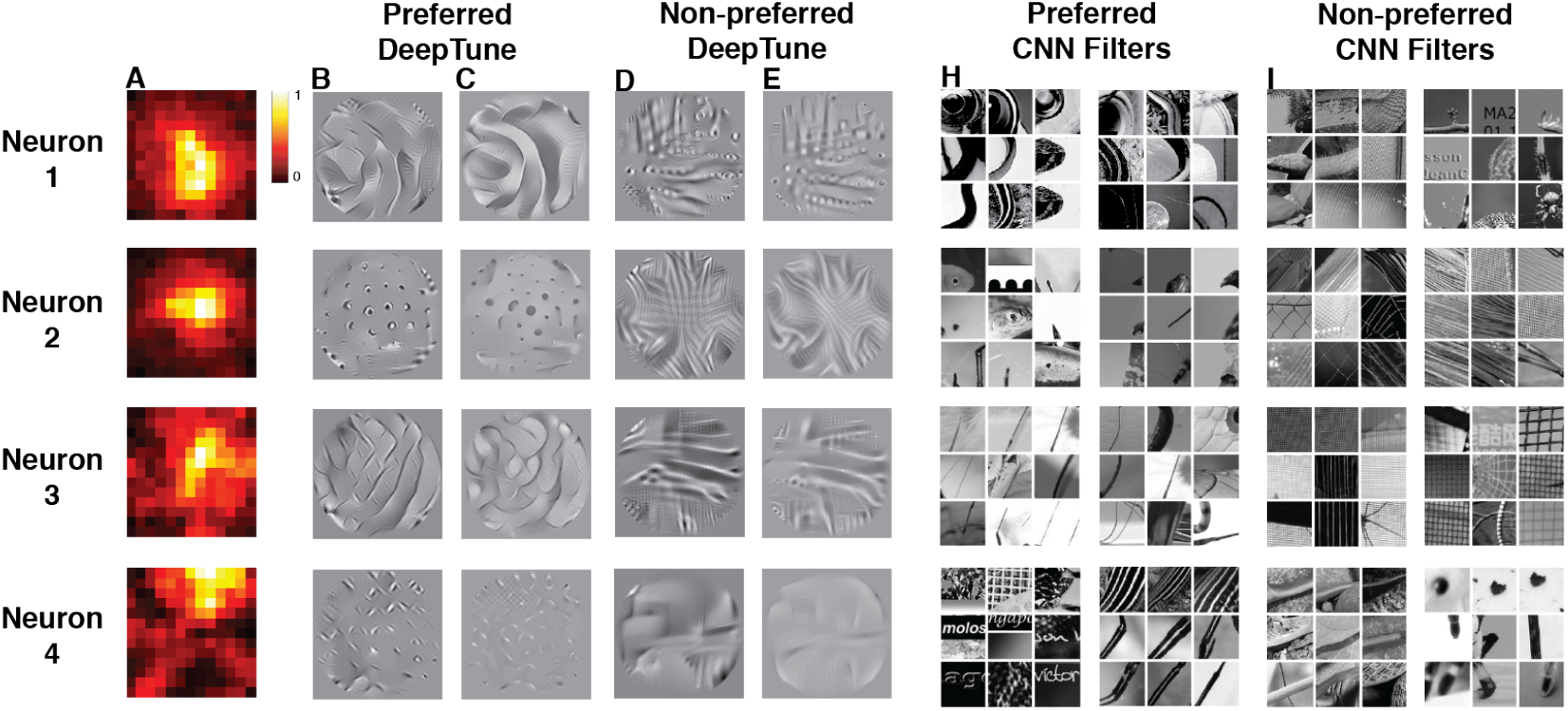
For Neurons 1-4, a comparison of excitatory and inhibitory DeepTune images, average regression weight maps and selected CNN features. **A.** Average regression weight map based on the AlexNet-Layer2 model. For each neuron, the average regression weight map also exhibits stable patterns across models (see *Stability of Analysis, Figure S11*) and it highlights the receptive field of a neuron. **B.** Excitatory DeepTune images from the AlexNet-Layer2 Model. Neuron 1 is tuned to the curvature-contour shapes with edges forming an approximately ninety-degree angle. Neuron 2 is selective for blob-like patterns and textures. A DeepTune image for Neuron 3 shows selectivity to curvature patterns with a strong diagonal line preference. Neuron 4 is tuned to corner-like shapes with edges forming ninety-degree angles. The rest of the 17 models show consistent patterns as shown in other DeepTune images (see *SI Stability of Analysis, Figure S8*). **C.** Excitatory consensus DeepTune images based on all 18 models. **D.** Inhibitatory DeepTune images from the AlexNet-Layer2 Model. **E.** Inhibitory consensus DeepTune images based on all 18 models. **H.** Top two excitatory CNN filters based on the filter importance index. To visualize a convolutional filter from a CNN, the 9 top image patches are presented from the ImageNet training set that have the highest filter responses. These 9 top image patches are representative of what this convolutional filter is computing [51, 50]. The top two selected CNN filters support the findings based on DeepTune images. For example, Neuron 1 is tuned for curved-contour patterns according to DeepTune images and its top CNN filters are those that activate on curvatures of similar shapes. Neuron 2 is selective for blob patterns and the top CNN filters activate respectively on blob pattern or pieces of a blob pattern. **I.** Top two inhibitory CNN filters based on the filter importance index.

The second question is: which types of convolutional filters contribute the most to the prediction performance? To address this question, we quantify the importance of each convolutional filter by *ℓ*_2_ pooling of the regression weights for a convolutional filter across spatial locations. Formally, for each neuron, the filter importance *I_m_* of *m*-th convolutional filter is defined as follows,

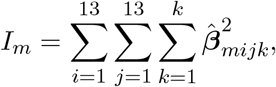

where *β̂*_*mijk*_ is defined as before. This filter importance index provides an independent view of neuron shape tuning through the most and the least important filters. To interpret the filter importance, a visualization of each convolutional filter in CNN is required. To this end, we adopt the filter visualization technique introduced by [51]. For each filter, we show the 9 top image patches from the ImageNet training set that have the highest filter responses (see *SI Methods* for visualization of AlexNet filters). These 9 top image patches are representative of what this convolutional filter is computing [51, 50]. Taking Neuron 1 as an example, Figure 4-F and G show the top and bottom two filters among 256 filters in AlexNet-Layer2 model ranked by the filter importance index, *I_m_*.

For each neuron, we observe that the top two filters capture essential image components corresponding to the tuning patterns shown in the DeepTune images. These tuning patterns are long curvatures for Neuron 1, blob-like patterns for Neuron 2, diagonal lines for Neuron 3, and corner-like shapes for Neuron 4. Comparing to the DeepTune images (Figure 4-B-C-D-E), the most important and least important CNN-features (Figure 4-C-H-I) provide an alternative interpretation of the excitatory and inhibitory tuning property of V4 neurons, respectively. Figure 4 shows that these two views (*I_m_* based and DeepTune) are visually consistent.

## 5 The wide variety of shape and texture tuning in V4

So far we have demonstrated that V4 neurons can be selective to both shapes (e.g. contour or curvature patterns) and textures. The finding that V4 are tuned to both shapes and textures agrees with previous studies using synthetic stimuli: on the one hand, V4 neurons are shown to be tuned to orientation and spatial frequency of edges and linear sinusoidal gratings [9], non-Cartesian gratings [10, 11] and curvature of contours [27, 28]; on the other hand, V4 is found to play a major role in processing textural information [23, 2, 26]. In order to further understand area V4 as a population of neurons, we use DeepTune as a new tool to investigate proportions of V4 neurons that are tuned to shapes, to textures, and to other patterns of stimuli.

Based on visual inspections of their consensus DeepTune images, we manually clustered our 71 neurons into five categories: two texture categories (blob-like and corner-like patterns), one for curved contours, one for lines and a final category for complex patterns. Figure 5-A is the count histogram of these five categories and Figure 5-B displays DeepTune images for three example neurons in each category with high correlation coefficients (¿ 0.4). This visualization again confirms that both texture-tuned and contour-tuned neurons are present in area V4. In fact, among the 71 neurons considered in this study, about 40% of them are selective to textures and 30% of them prefer contour shapes. A finer manual categorization shows that among the ones selective to textures, half are tuned to blob-like patterns and the other half prefer corner-like patterns. Contour-selective neurons show preferences to either curvatures or straight lines like some typical V1 neurons but with larger receptive fields. The number of neurons selective to curved contours is twice of that selective to straight lines. We have also included in the last category the neurons tuned for complex patterns that are hard to describe in language and do not fall into previous categories. By displaying neuron tuning in a concrete and naturalistic manner, the DeepTune images extends the results in previous studies on V4 neuron selectivities [11, 17].

**Figure 5:**
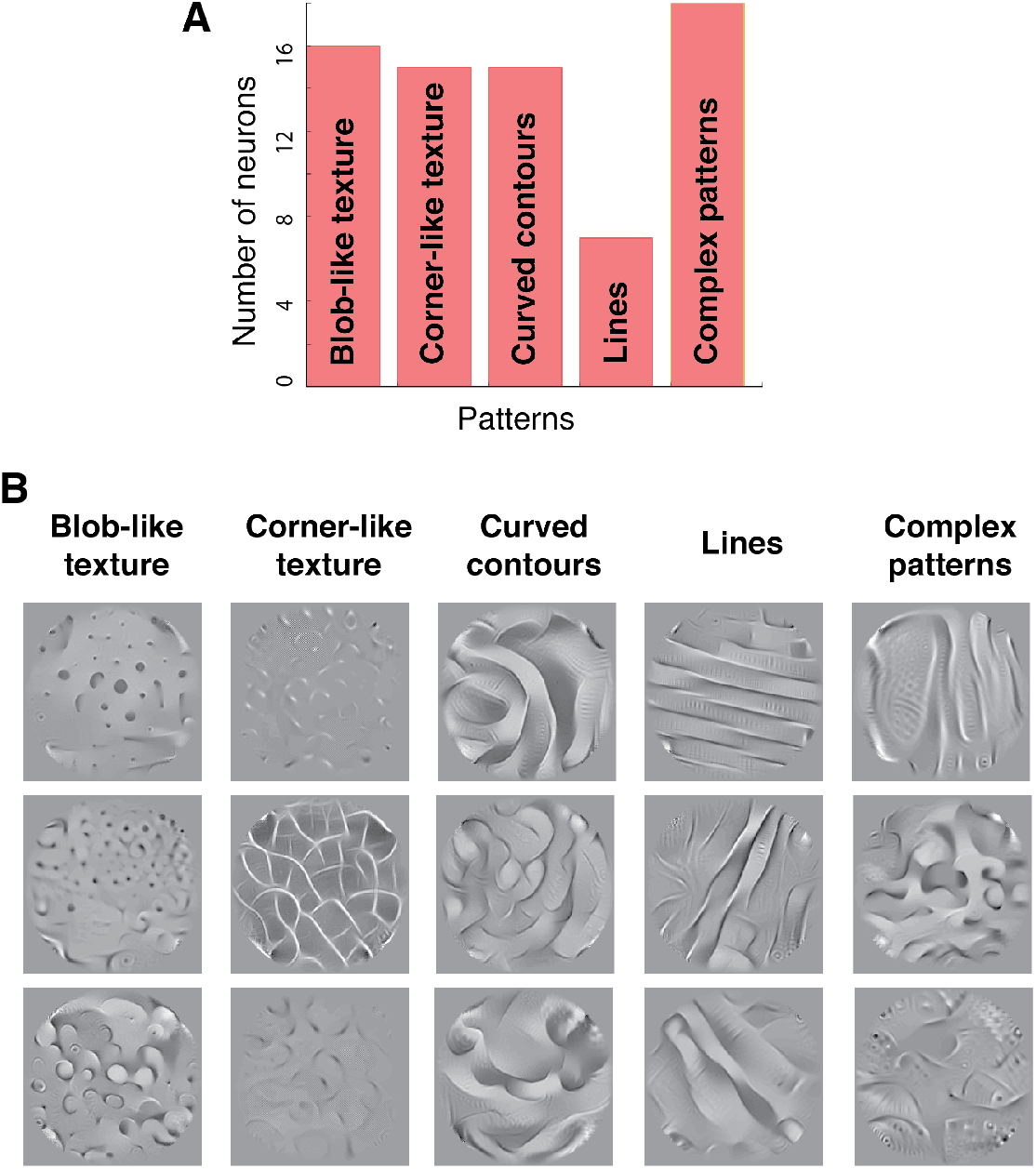
Diversity of tuning among 71 V4 neurons. **A.** Neurons are manually categorized into five categories based on their DeepTune images. More than 40% of the neurons are selective to texture, half of which prefer blob-like textures and the other half prefer corner-like textures. About 30% of the neurons exhibit contour patterns, both curvature and straight lines. Neurons selective to curvatures are twice as the ones selective to straight lines. The rest of the neurons have selectivities to visually complex patterns. **B.** Examples of consensus DeepTune images for three neurons from each of the five categories.

## 6 V4 curvature tuning to a full range of separation angles

It is suggested by Roe et al. [30] that diverse curvature tuning in V4 provides an efficient way to encode shapes. However, it is not yet clear that how different types of curvature tunings are distributed in the V4 population. Previously, artificial curvature stimuli have been used to probe the different angle tuning properties in area V4 [27, 28]. These stimuli are constructed by joining two oriented line segments in a sharp corner or curve. These studies highlight the presence of bimodal orientation tuning with various separation angles. The preferred separation angle is defined in [28, 8] as the angle between the two most preferred oriented line segments passing through the center. The SRF analysis [8] also confirm bimodal orientation tuning in V4 by showing the presence of neurons tuned sharp corners. As for the distribution of different angles, Carson et al. [6] observed that not all curvatures are equally represented. They use sparse modeling of object coding to show that the strong representation of acute curvatures across the neural population. In this section, we investigate whether DeepTune images can concretize previous discoveries and provide visualization of V4 neurons tuned to different separation angles.

By visually inspecting the consensus DeepTune images of 71 V4 neurons, we first identified the 38 neurons that are tuned to curved contours, corner-like shapes and lines. Then we manually clustered these 38 neurons into four categories based on their separation angles of their curves (45°, 90°, 135°, 180°). Figure 6-A shows a count histogram of (excitatory) separation angle of the 71 V4 neurons. We observe that there is a strong presence of neurons with curvature tuning at less or equal to 90° separation angles (18 out of 71 neurons). Another 15 neurons are selective to blob-like textures that does not correspond to any particular angle. There are 18 neurons that are not selective to any clear angle or blob-like patterns.

**Figure 6:**
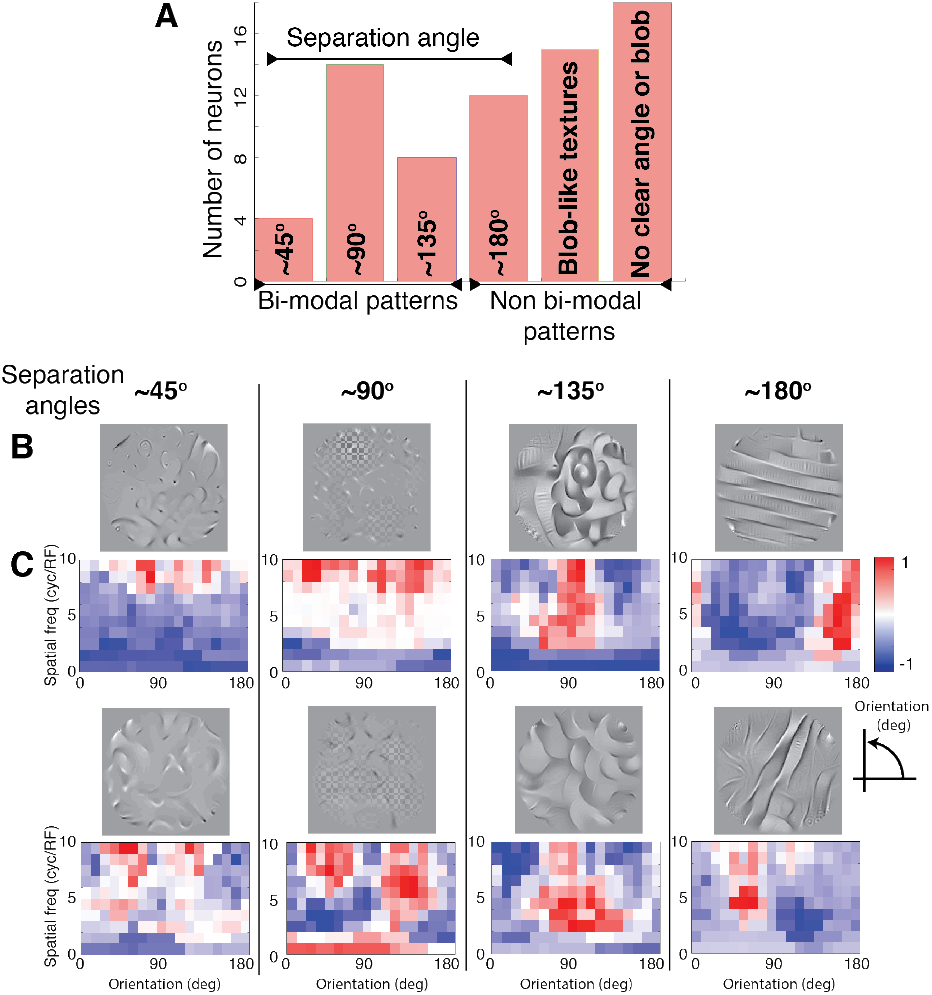
Categorization of V4 neurons based on their separation angles. **A.** Neurons are manually categorized into six groups. The first four groups contains neurons tuned to patterns with separation angles of 45°, 90°, 135°, and 180°. These patterns are either contours or textures. About 20% out of 71 neurons are tuned to patterns with separation angles of 90°. Another 20% of the neurons are selective to blob-like textures that do not correspond to any particular angle. The rest of neurons are not selective to any clear angle or blob-like patterns. **B.** The consensus DeepTune images for two example neurons in each of the first four categories and **C.** The corresponding spectral receptive field (SRF) (David et al [8]) visualization. The orientation tuning obtained via SRFs and consensus DeepTune images are consistent. while SRF predicts a neuron has tuning for a particular angle through Fourier analysis, the consensus DeepTune images offer concrete visualization of these tunings. For example, for the bottom left neuron, both our method and SRF show an orientation tunings of about 70° and 120°.

To further support the separation angles for V4 neurons identified by looking at DeepTune images, we perform spectral receptive field (SRF) analysis [8] on our data and compare the angles identified by both analyses. In Figure 6-B and C, for each neuron, we display in one column the consensus DeepTune image and the SRF plot as in David et al. [8]. The horizontal axes of the SRF show the orientation tuning of each neuron, with preferred component in red. In the SRF plot, according to [8], the separation angle corresponds to the difference between the top two orientation tuning peaks. We observe that the separation angle from the SRF plot are consistent with the ones from the DeepTune images. For example, for the bottom left neuron, both DeepTune and SRF show two orientation tuning peaks at about 70° and 120°. To summarize, the diversity of excitatory curved-contour patterns in fact matches the previous neurophysiological observations in V4 [28, 6, 24]. Furthermore, our DeepTune images offer a concrete visualization of the bimodal orientation tuning properties of many V4 neurons, refining earlier analysis.

## 7 Suppressive tuning discovery via inhibitory DeepTune

It is well known that V4 neurons have surround suppressive mechanisms [9, 33, 18] just like many other visual cortical areas [14, 1]. Besides, recent study by Willmore et al. [45] found evidences for the presence of strong suppressive tuning to specific features in about half of the neurons in area V2. In addition, they show that this type of suppressive tuning is not caused merely by surround suppression and is not present in area V1. In this section, we investigate whether such strong suppressive tuning is also present in area V4.

To study the suppressive tuning in the area V4, we fit the Berkeley Wavelet Transform (BWT) model [45] to our data. The BWT-based model provides a nonlinear spatio-temporal receptive field (STRF) for each neuron. We adopt the excitation index (EI) introduced in [45] as:

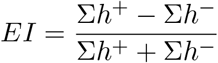

where *h*^+^ and *h^−^* are positive and negative weights respectively assigned to the wavelets in each STRF.

The BWT-based model has an average prediction correlation coefficient 0.33 for the 71 V4 neurons in the holdout test set. It is about 0.09 lower than the worst among 18 CNN-based models. While this model does not fully explain the non-linear property of V4 neurons, its accuracy is comparable to that of the same BWT model for V2 neurons (average correlation coefficient of 0.30) [45]. Figure 7-A shows the histogram of excitation index for 71 V4 neurons. 41% of the neurons in V4 show suppressive tuning. The median of the excitation index for V4 neurons is 0.10. While the portion of neurons with suppressive tuning is 9% lower compared to that in V2, it is 29% higher than that in cortical area V1 [45].

**Figure 7:**
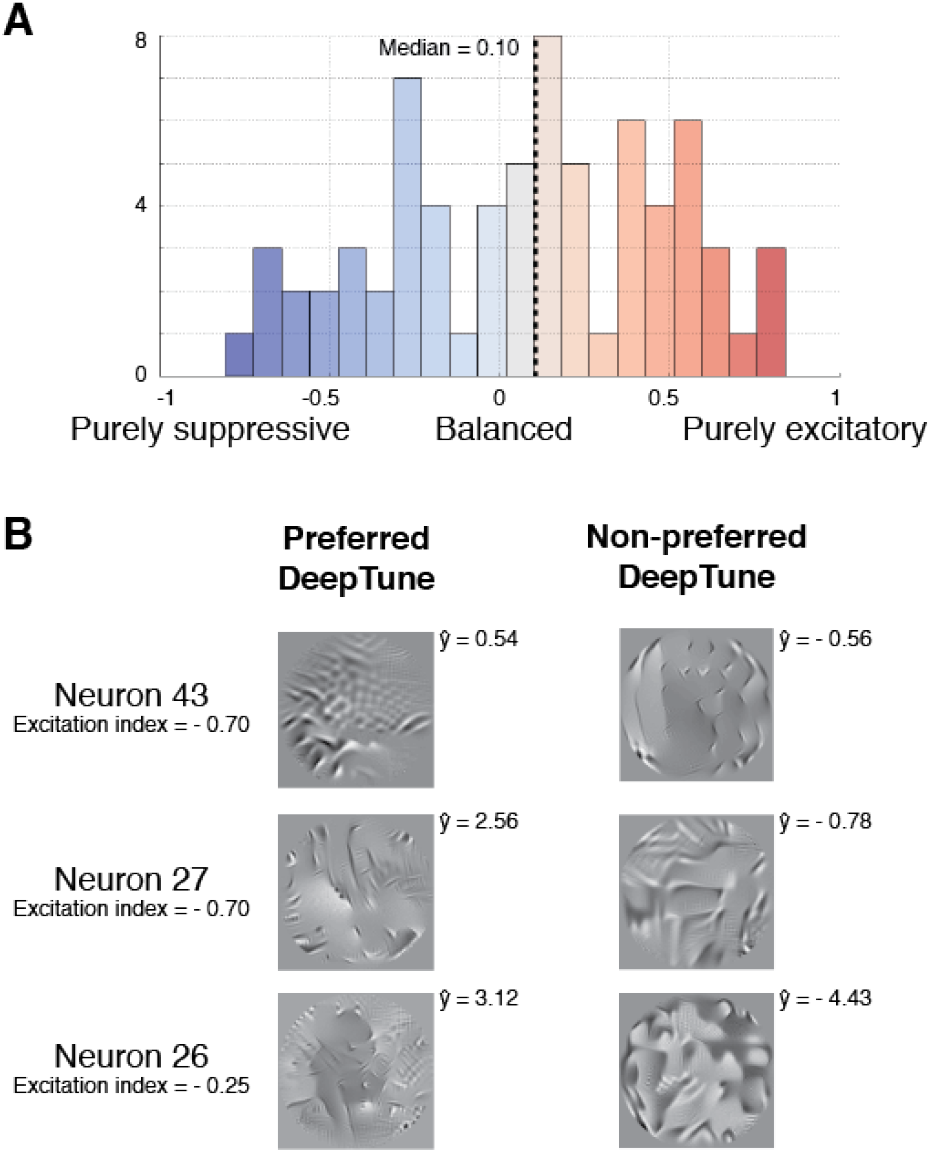
Neurons in the primate cortical area V4 exhibit suppressive tuning. **A.** Histogram of BWT excitation index for 71 V4 neurons. 41% of the neurons show strong suppressive tuning. The median of excitation index for V4 neurons is 0.10. **B.** The excitatory and inhibitory DeepTune images for three neurons identified as suppressive by the BWT model. The neuron excitation index and response of the model to each DeepTune image is illustrated in the same panel. The neurons with suppressive tuning have much clearer suppressive DeepTune images than those without. *ŷ* is the predicted model response obtained by feeding the DeepTune image through AlexNet-Layer2 model.

Figure 7-B presents the excitatory and inhibitory consensus DeepTune images for three neurons identified as suppressive neurons according to the BWT model (on the left side of the histogram). The corresponding excitation indexes are shown below the neuron names. Recall that the excitatory DeepTune images are obtained via maximizing the model response (with appropriate regularization), while the inhibitory ones are obtained via minimizing the model response (with appropriate regularization). The neuron excitation index and response of the model to each DeepTune image are shown in the same panel. For example, *ŷ* = 0.54 means that the model predicted a firing rate of 0.54 spikes per sampling period (16.7ms). The DeepTune images provide a concrete visualization of the suppressive tuning in V4: The excitatory DeepTune images of these neurons are weak and/or blurry, while the inhibitory DeepTune images show sharper patterns. In the case of Neuron 43, while the excitatory DeepTune has blurry patterns, the inhibitory DeepTune exhibits a clear tuning to ninety-degree corner shapes in the right hand side of visual field. This means that a ninety-degree corner shape is likely to drive this neuron firing rates close to zero. Moreover, looking at the other inhibitory DeepTune images, both of neurons 27 and 26 have strong suppressive tuning to complex shapes with mid-range frequency.

## 8 Discussion

Prior work has demonstrated the power of deep CNNs in building accurate predictive models of neural responses in V4 [48, 4]. In this work, we have similarly demonstrated that pre-trained CNNs give state-of-the-art results in modeling V4 neuron responses to natural images. We additionally have presented the DeepTune framework for eliciting stable visualizations and interpretations of these models. The generated visualizations are stable over modeling choices and randomness in the model training procedure.

### 8.1 Flexible visualization of optimal stimuli

The idea of computationally optimizing input stimulus to discover neuron tuning properties dates back to Carlson et al. [6]. The evolutionary sampling method was used to optimize for the stimulus that causes the highest number of spikes. This work greatly expanded the search space of tuning patterns compared to previous methods that were based on handcrafted stimuli [27, 10]. However, the evolutionary sampling method in [6] is constrained on limited concatenated Bezier splines. It can generate spline-based contours easily, but has difficulty for generating fine-scale texture stimuli. Our DeepTune images are generated from a regularized optimization directly over the input pixel values, and hence have an even larger search space that allows for more complex and naturalistic tuning patterns.

The resulting DeepTune population analysis demonstrates that V4 neurons are tuned to a huge variety of shapes as well as textures in different orientations. It also reveals that the tuning properties of many V4 neurons cannot be explained by simple edge and corner patterns. We see in Figure 7, for example, that even the stable part of the DeepTune images is difficult to describe in such simple terms. This suggests that tuning in area V4 is much more complex than that of V1 and than what can be described by handcrafted grating stimuli. Studies based on synthetic stimuli [10, 11, 27] may lack the expressive power to represent shapes of many V4 neuron receptive fields. Predictive modeling approaches as SRF [8] may not be sufficient to capture the complex tuning properties either. It provides only summary statistics such as spatial frequency and orientation about the receptive fields.

### 8.2 Distinctions in curvature selection revealed by DeepTune

Examining the DeepTune images of Neuron 3 and 4 in Figure 4, we see that both neurons are tuned to curvatures with similar edge orientations (two edge directions with a separation angle of ninety degrees). However, they have very distinct shape tuning properties apart from the orientation tuning summary statistics. Neuron 3 prefers a curvature-contour pattern with a ninety-degree angle and long edges. Neuron 4 prefers a corner-like repeated texture. This agrees with the study by Nandy et al. [24]. It is suggested that the curvature selection of V4 neurons could arise for two reasons: systematic variation in fine-scale orientation tuning across spatial locations (like Neuron 3), and local tuning heterogeneity (like Neuron 4). Note that this type of refined result would be difficult to obtain via methods based on global Fourier analysis such as spectral receptive field (SRF) [46, 8]. The 2D Fourier transform is spatial translation-invariant, meaning it is difficult to distinguish between Neuron 3 and Neuron 4 via SRF analysis.

### 8.3 DeepTune to accelerate neurophysiology experiments

The DeepTune images for each V4 neuron are concrete and naturalistic. They are visually very similar to many input image stimuli. In other words, the DeepTune images are ready to be fed back to neurons as stimuli for confirmation or refutation of their characterizations of tuning properties in a closed experimental loop. Consequently, DeepTune images hold the promise to speed up the efficiency of data collection in V4 and other brain areas.

## 9 Methods

### Electrophysiology

Extracellular recordings were made from well isolated neurons in parafoveal areas V4 (71 neural sites) of three awake, behaving male rhesus macaques (Macaca mulatta). Surgical procedures were conducted under appropriate anesthesia using standard sterile techniques [43]. Areas V4 were located by exterior cranial landmarks and/or direct visualization of the lunate sulcus, and location was confirmed by comparing receptive field properties and response latencies to those reported previously [12, 34]. All procedures were done in accordance with National Institutes of Health guidelines. See *SI Data Collection* for additional details.

### Software Packages

The regularized linear regression analysis is performed using the SPAMS package [22]. The neural network feature extraction and the DeepTune framework are implemented using the Caffe package [15]. The pre-trained neural network architectures are from the Model Zoo of the Caffe package. See *SI Methods* for additional details.

## Acknowledgements

This research is supported in part by ONR Grant N00014-16-2664, NSF Grants DMS1613002 and IIS 1741340, and the Center for Science of Information (CSoI), a US NSF Science and Technology Center, under grant agreement CCF-0939370. We would like to thank Bruno Olshausen and Friedrich Sommer for fruitful discussions, Julien Mairal for fruitful discussions and sharing SPAMS code snippets, and Anwar Nunez-Elizalde for helpful comments on the presentation of the paper.

## Supporting Information for “The DeepTune framework for modeling and characterizing neurons in visual cortex area V4”

### A Data Collection

Extracellular recordings were made from well isolated neurons in parafoveal areas V4 (71 neurons) of three awake, behaving male rhesus macaques (Macaca mulatta). This dataset has been previously used to study the sparseness of neural codes in the area V4 [44]. Surgical procedures are thus identical to those in [44]. We restate the procedures here for completeness. Surgical procedures were conducted under appropriate anesthesia using standard sterile techniques [43]. Areas V4 were located by exterior cranial landmarks and/or direct visualization of the lunate sulcus, and location was confirmed by comparing receptive field properties and response latencies to those reported previously [12, 34].

All procedures were approved by the Animal Care and Use Committee of the University of California, Berkeley, and were conducted in strict accordance with good practice as defined by the Office of Laboratory Animal Care at UC Berkeley, the National Institutes of Health, the Society for Neuroscience, and the American Association for Laboratory Animal Science.

During recording, the animals performed a fixation task for a liquid reward. Eye position was monitored with an infrared eye tracker (500 Hz; Eyelink II; SR Research) and trials during which eye position deviated *>* 0.5° from the fixation spot were excluded from our analysis. The standard deviation of the fixational eye movements was typically 0.1°. Activity was recorded using tungsten electrodes (FHC), and amplified and neural signals were isolated using a spike sorter (Plexon).

Experiments were controlled and stimuli generated using custom behavioral/stimulus display software (PyPE) running on a Linux-based PC. Stimuli were displayed on a 21 inch Trinitron monitor (Sony) capable of displaying luminances up to 500*Cd/m*^2^. The luminance nonlinearity (gamma) of the monitor was calibrated and corrected in software to provide a linear luminance response.

In the main experiment, each neuron was probed with a rapidly changing sequence of natural images. The images were circular patches of grayscale digital photographs from a commercial digital library (Corel). Patches were chosen by an automated algorithm that selected them at random but favored patches with high contrast [to reduce the frequency of blank stimuli (e.g., patches of sky)]. All patches were adjusted with a gamma nonlinearity of 2.2, to give an appropriate luminance profile on our linearized display. The outer edges of the patches (10% of the radius) were blended smoothly into the neutral gray background, whose luminance was chosen to match the mean luminance of the image sequence.

Random images were then concatenated into long sequences so that each 16.7 ms frame contained a random image patch from the library. All images were centered on the classical receptive field (CRF) and patch size was adjusted to be two to four times the CRF diameter. CRF diameters ranged from 0.5 to 10.4° (median, 2.2°), and eccentricities ranged from 0.1 to 49° (median, 3.1°). The entire sequence was broken into 3–5 s segments, and one segment was presented on each fixation trial. To avoid transient trial onset effects, the first 196 ms of data acquired on each trial were discarded before analysis.

The training dataset of a neuron consists of 4, 000 *−* 12, 000 natural images (See Figure 8 for a random sample). Spike count was measured at 60 Hz, resulting in two measurements per image. For the holdout dataset, 300 images were shown for each neuron, distinct from the images shown for the training dataset. The sequence of test images was repeated; on average, each image in the test set was shown 9.3 times. The resulting spike counts were averaged to provide a lower-variance estimate of the expected spike count; repeats also allowed for estimation of the amount variance in the neuron explainable by the stimulus image (signalto-noise).

**Figure 8:**
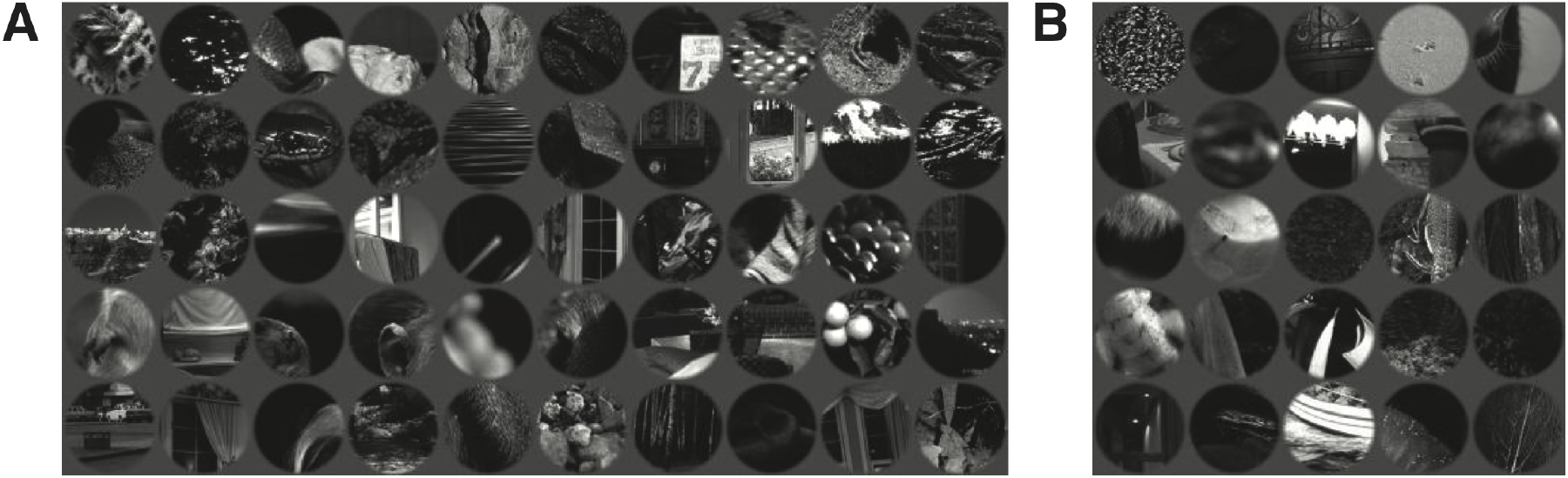
Sample of Images from training and holdout datasets. A. 50 images sampled from training dataset of 4000 images of one neuron. B. 25 images sampled from holdout dataset of 300 images of one neuron.

### B Methods

In this section, we discuss the methods we used to model neurons and the metrics to characterize our models’ performance compared to measured neuron activity.

#### Single Neuron Modeling and Metrics

As described in the main text, we use transfer learning framework to analyze sinlge V4 Neuron input-response data: We first extract convolutional neural networks (CNN) features and then use as predictors in a linear regression method to predict spike-rates as the response. The CNNs are pretrained on large scale image classification dataset ImageNet [32]. The linear model learned by regularized linear regression is trained on our data.

As a measure of the prediction performance of our model, the correlation between the expected spike count predicted by the model and the actual average spike count on the holdout set is computed.

*Explainable variance* captured by the model is another relevant metric for predction performance in the neuroscience litterature [29, 35]. This metric attempts to control for differences in noise levels between experimental setups, individual neurons, and brain regions. We estimate explainable variance using the repeat presentations of images in the test set.

#### Convolutional Neural Networks (CNN)

Deep convolutional neural networks are a successful tool to analyze big data problems and are therefore are being actively studied for a vast variety of applications especially in machine learning [20, 19, 36].

Convolutional networks are basically neural networks with several layers and a specialized connectivity structure. The purpose is to extract features of the scene in multiple layers. It has been shown that higher layers compute more global features than lower layers, so that the hierarchical structure provides a better overall quality of features [51]. The proposed architecture for several layers of network varies in different applications but it usually consists of three general types: convolutional layers, pooling layers and fully-connected layers.

*Convolutional layers* select a window of previous layer’s output and convolve it with a set of filters. Dependencies are local in this structure. The coefficients of these filters are tunable weights of our network and their final value will be specified in the training procedure. As an example, considering images as the input of our network, each filter is a rectangular grid which will be convolved with specific patch of previous layer. A non-linear function will be used to specify the output of the neuron as in traditional neural networks. Equation 1 specifies this relationship between output of different layers for two-dimensional configuration.

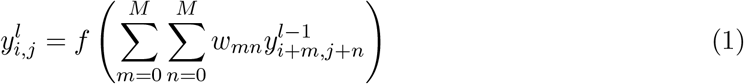

where *i* and *j*’s indicates possible spatial location at layer *l*, 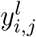 is the output of each neuron in layer *l*, *w_mn_* is the filter weights at location (*m, n*) of layer *l −* 1 and *f* is the non-linear function.

*Pooling layers* could be utilized after each convolutional layer. It simply performs a spatial pooling over patches of previous layer. These patches could be overlapping or non-overlapping. The output of pooling layer for each patch is a single value which in most of the cases is maximum value of the patch. Pooling could be useful to reduce the feature dimension as well as increase the invariance of the features for small transformation. It also helps to increase the size of receptive field for each feature value.

After several convolutional and pooling layers aimed at grasping the low-level and high-level features, a few *fully-connected layers* are used as the final stages of the network. These layers are essential for specific application of the network such as classification or prediction.

Figure 9 shows the neural network architecture of the AlexNet model [19]. It consists of five convolutional layers, three pooling layers inbetween and two fully connected layers. Our analysis is carried out on all the seven layers shown in the figures. Layers L2, L3 and L4 are of the main focus. In particular, the output feature at L2 is of size 256 *×* 13 *×* 13, where 256 indicates the number of types of filters applied at Layer L2, 13 *×* 13 indicates that the features are extracted on a spatially-equally-spaced 13 *×* 13 overlapping grid of the original image. Similarly the output features at layers L3 and L4 are of size 384 *×* 13 *×* 13 and 384 *×* 13 *×* 13.

**Figure 9:**
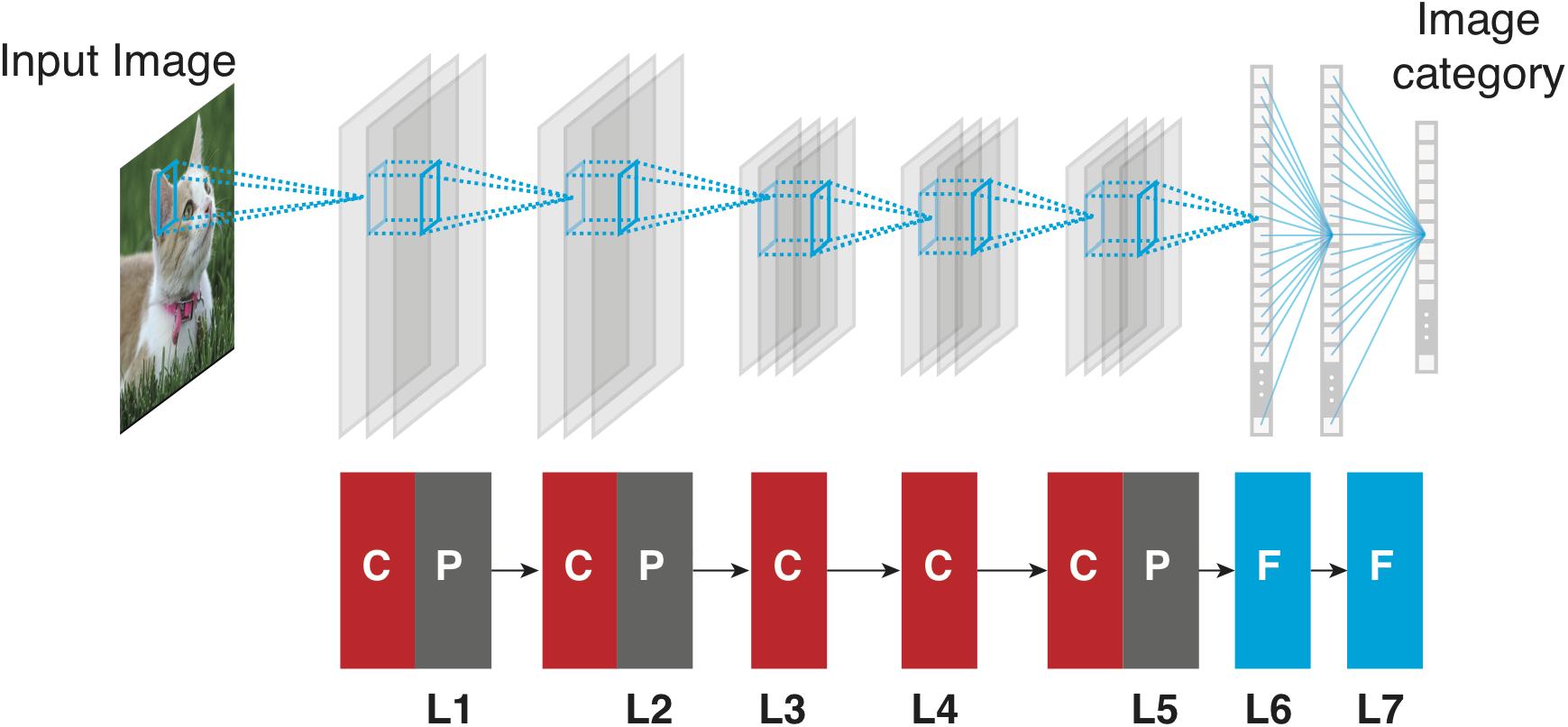
Architecture of the AlexNet model [19]. Red box indicates convolutional layer, gray box indicates pooling layer and blue box indicates fully connected layer.

We also used GoogLeNet [39] and VGG [38] in our analysis. The architectures and the mechanism of these models are beyond the scope. We refer the readers to the original paper for a detailed understanding. The CNN feature extraction pipeline is done using the Caffe [15] package and the model files provided within.

#### Regression Methods

As described in the main text, our predictive model for a single neuron response takes the following form

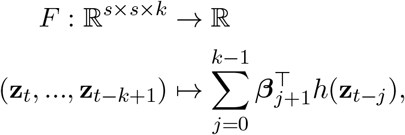

where (*β*_1_, …, *β_k_*) ∈ ℝ^*d×k*^ are the regression parameters to be determined and *h* is the fixed CNN feature transform.

To perform the regression analysis, we solve the following regularized linear regression problem

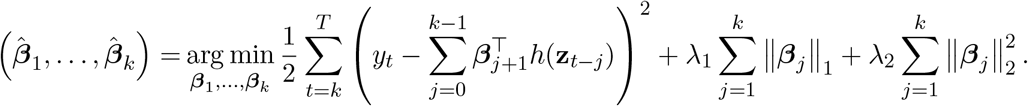

Taking the AlexNet Layer 2 model as an example, the Layer 2 feature is of dimension *d* = 256 *×* 13 *×* 13. Taking into account the time lags, the weight matrix (*β̂* _1_, …, *β̂* _*k*_) is of dimension 256 *×* 13 *×* 13 *×* 9. This feature dimension is much larger than the sample size *T* = 8000. Regularization methods are needed to both improve prediction accuracy and provide better interpretation.

Either ridge regression (*ℓ*_2_ regularization) or LASSO [40] (*ℓ*_1_ regularization) will be suitable for this high dimensional regression problem. While ridge regression is commonly used in the neuroscience litterature, LASSO could provide better guarantees for feature selection [52] in theory. We find that both regression methods produce consistent prediction performance and DeepTune images in our analysis. A detailed comparison is discussed in Section C.

#### Visualization of CNN filters

In this subsection, we provide visualization of CNN filters. These visualizations show that CNN features encode much richer patterns that Gabor wavelets do. They support our finding that CNN based models perform better than simple Gabor wavelet based models in modelling V4 neurons. Because of the pooling operations, normalization operations, and non-linear activation functions in the CNNs, the CNN features are complex nonlinear functions of the raw input image. These CNN features are the outcome of learning from the large scale image dataset ImageNet, and are in general hard to explain via mathematical formula.

Inspired by the recent advances in CNN visualization [51, 50], we visualize the filters Layer 2, 3 and 4 of the AlexNet as follows: taking Layer 2 as an example, for each of the 256 types of filters, we exaustively search for nine image patches, from a dataset of one million image patches generated from ImageNet, that has the maximal output responses for the filter. The one million image patches are generated by randomly cropping images in ImageNet.

Figure 10 shows a subset of 256 types of filters in Layer 2 of AlexNet. We have manually clustered these filters in categories. We observe that other than encoding edge-shape patterns, Layer 2 of AlexNet also encodes a rich set of curvature patterns, contour-blob patterns as well as crossing patterns. These patterns could be very useful in building a predictive model for V4 neurons, because similar shape tuning properties of V4 neurons have been reported before [30].

**Figure 10:**
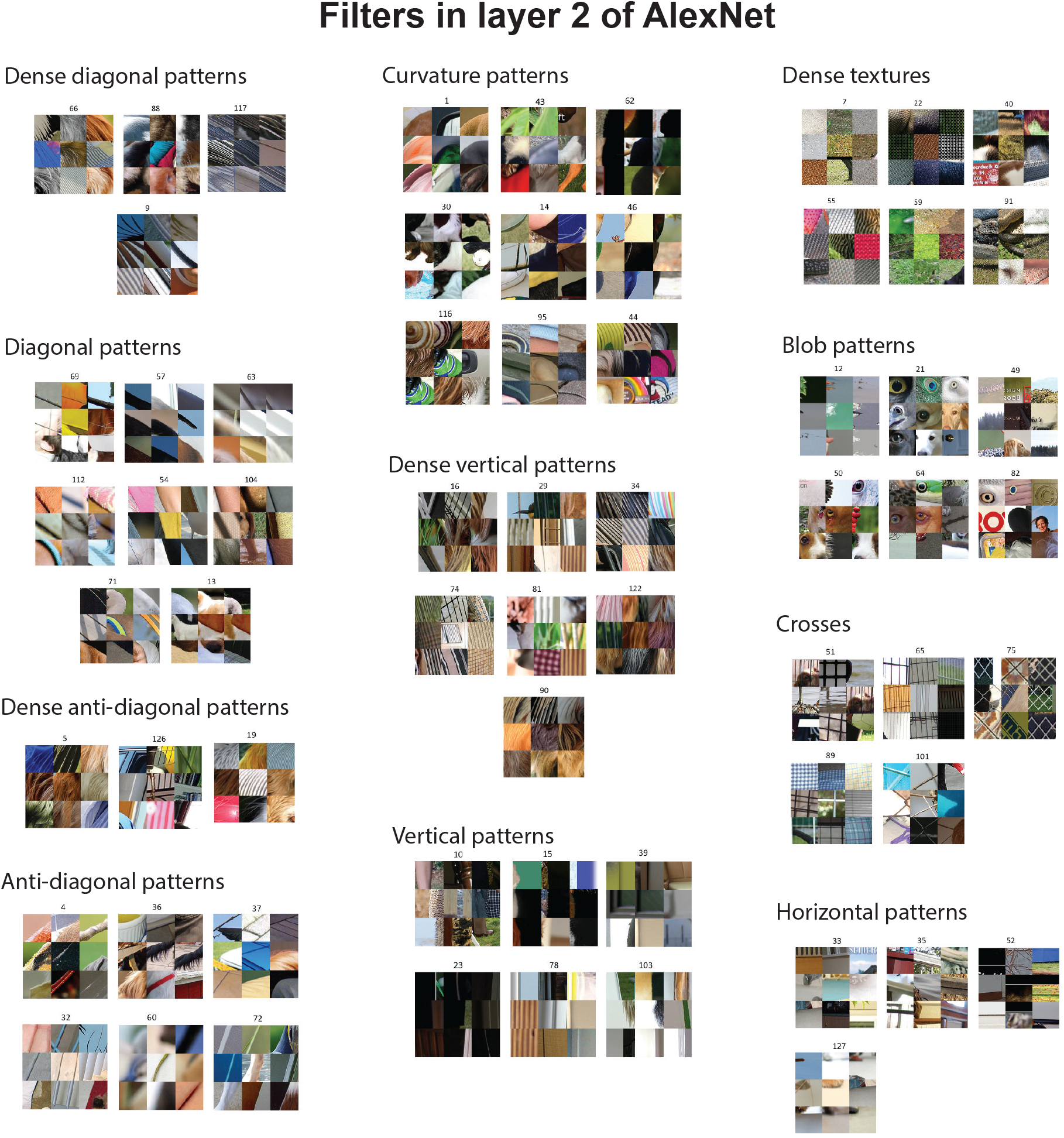
Subset of filters in Layer 2 of AlexNet. To visualize each filter, we have fed one million image to the CNN and visualized top nine image patches that activate that has the maximal output responses for the filter [51]. We have manually clustered filters into categories.

Similarly, Figure 11 and Figure 12 shows a subset filters in Layer 3 and Layer 4 of AlexNet. We observe that these filters encode even richer shape patterns. Some concrete patterns such as “dog head” and “birds” appear in Layer 3 filters. It has been shown that higher layers compute more global complex features than lower layers [51]. Unfortunately, the higher layer features become more specific to the classification task used to train AlexNet. It is not clear the higher layer features are as transferable as the lower layer features to other tasks [36].

**Figure 11:**
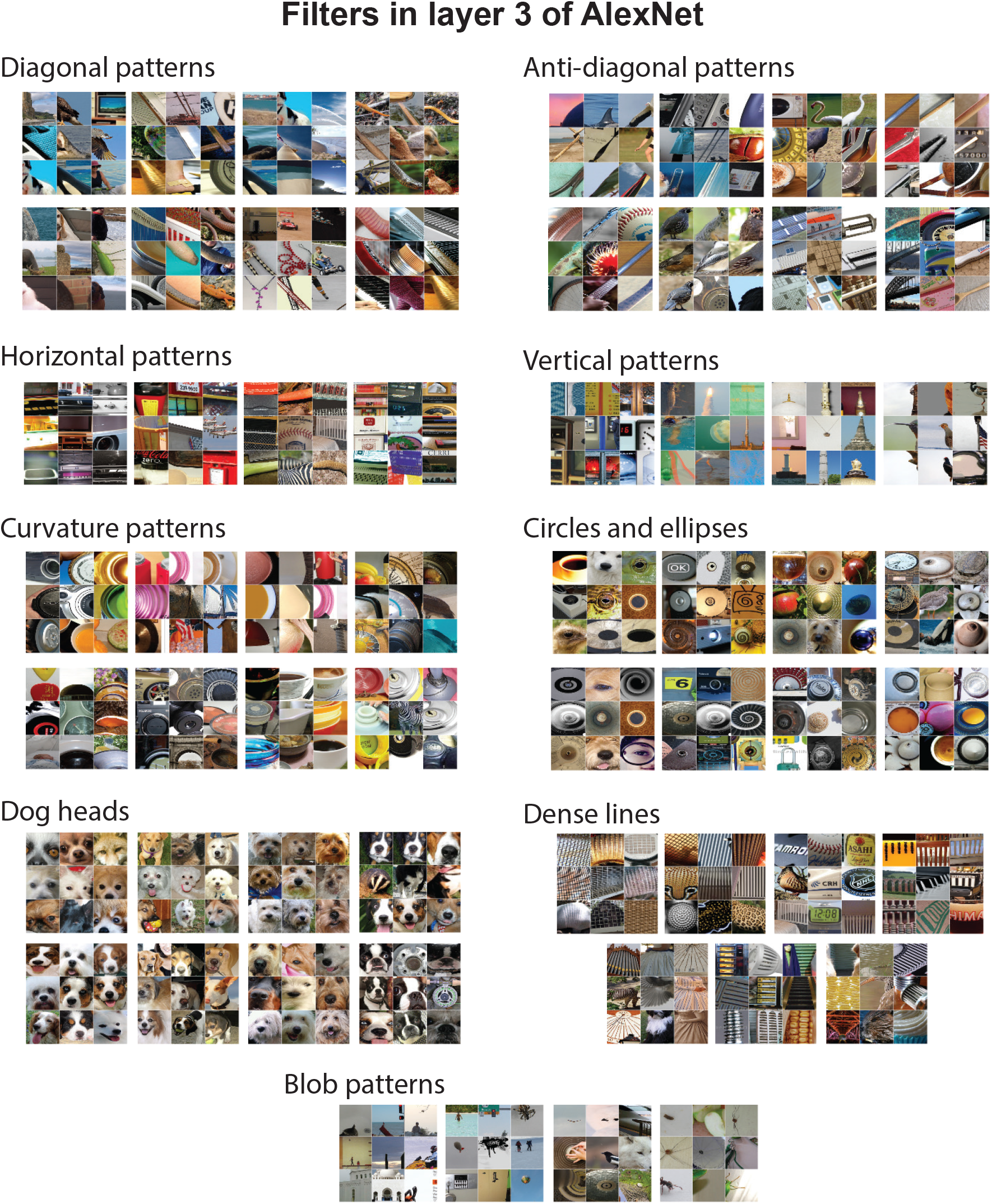
Subset of filters in Layer 3 of AlexNet. To visualize each filter, we have fed one million image to the CNN and visualized top nine image patches that activate that has the maximal output responses for the filter [51]. We have manually clustered filters into categories.

**Figure 12:**
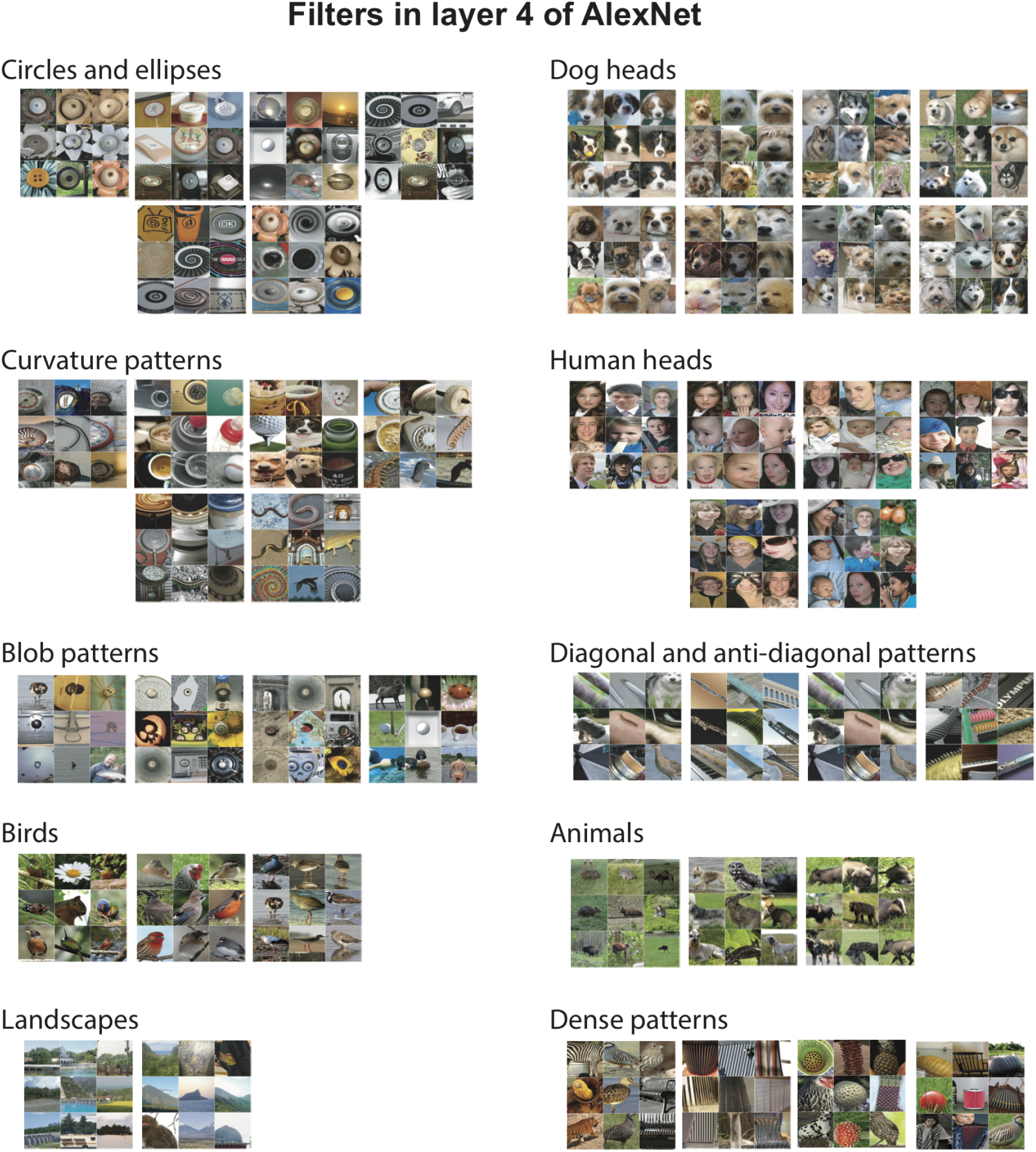
Subset of filters in Layer 4 of AlexNet. To visualize each filter, we have fed one million image to the CNN and visualized top nine image patches that activate that has the maximal output responses for the filter [51]. We have manually clustered filters into categories.

### C Stability of Analysis

In this section, we discuss the stability of our analysis for DeepTune images and model selected features.

#### Stability of DeepTune Images

Our main analysis is based on DeepTune Images. The CNN-based approach for interpretation is potentially biased because of the specific choice of architecture, parametrization and methods. In this section, we investigate the convergence and stability of DeepTune visualization to different perturbations. Additionally, we study the stability of selected CNN features and weight-maps.

#### Convergence

To visualize the DeepTune image optimization process, we use SuperHeat visualization package to plot the heatmap of the CNN feature activation map throughout the optimization process in Figure 13. There is a transition of the CNN feature activation map at about DeepTune iteration 8. After this iteration, the CNN feature activation map stabilizes. The inactive columns correspond to the color-selective features in AlexNet. Our stimulus is gray-scale, therefore, it is expected to observe weak selection for these filters.

**Figure 13:**
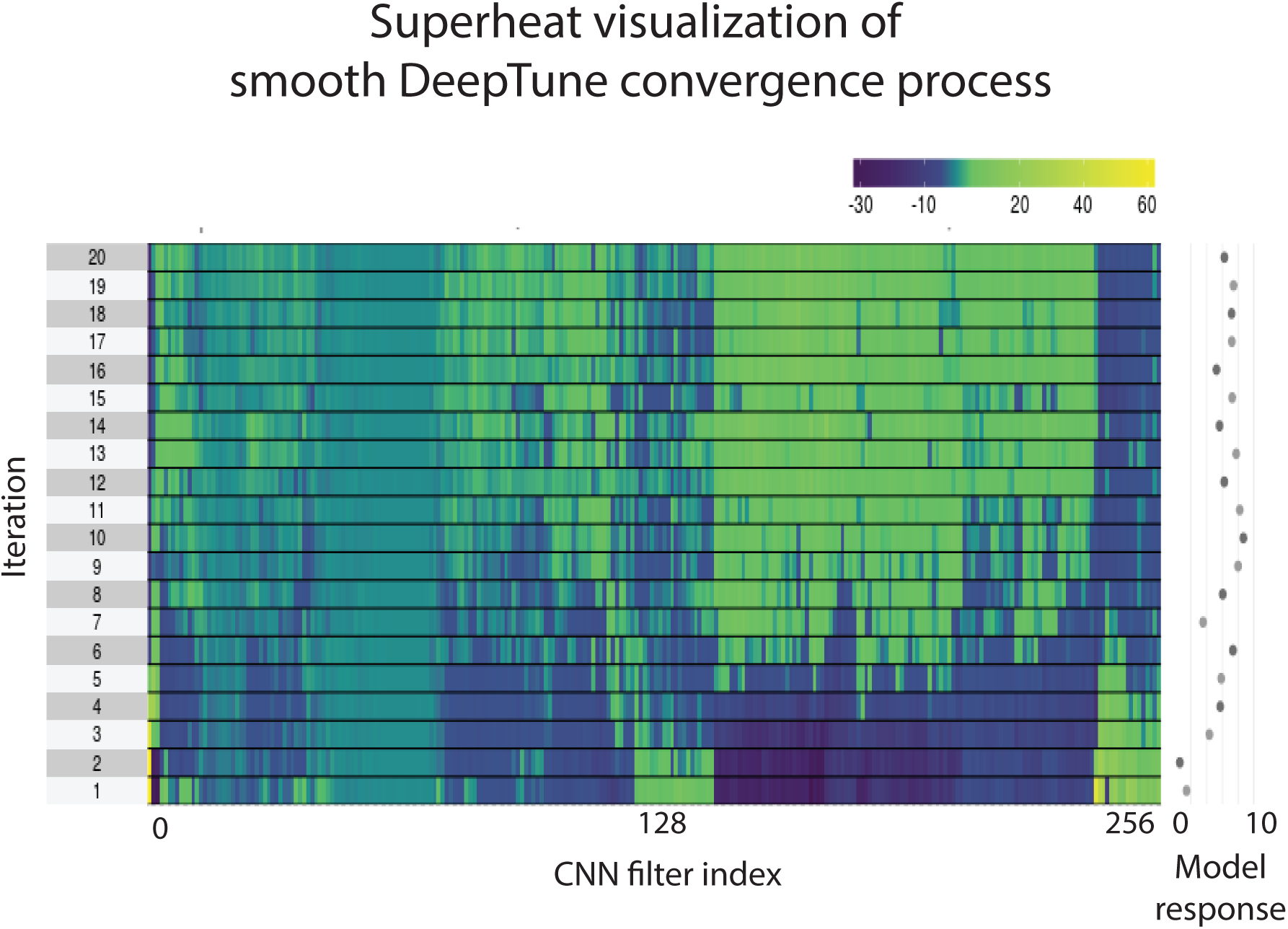
The DeepTune image optimization process. We use SuperHeat visualization package to plot the heatmap of the CNN feature activation map throughout the optimization process.

#### Stability across different initialization

A DeepTune image is the final result of an optimization process on an initial random image. To study the effect of random initialization on the final DeepTune image, we run the optimization process on 10 different random starting image for each neuron. Figure 14 shows 10 DeepTune images from these different initializations for five neurons. The patterns from 10 DeepTune images are visually similar. The average pair-wise correlation coefficient between 10 images is 0.97 for neuron 1. For other neurons, this value is not less than 0.94.

**Figure 14:**
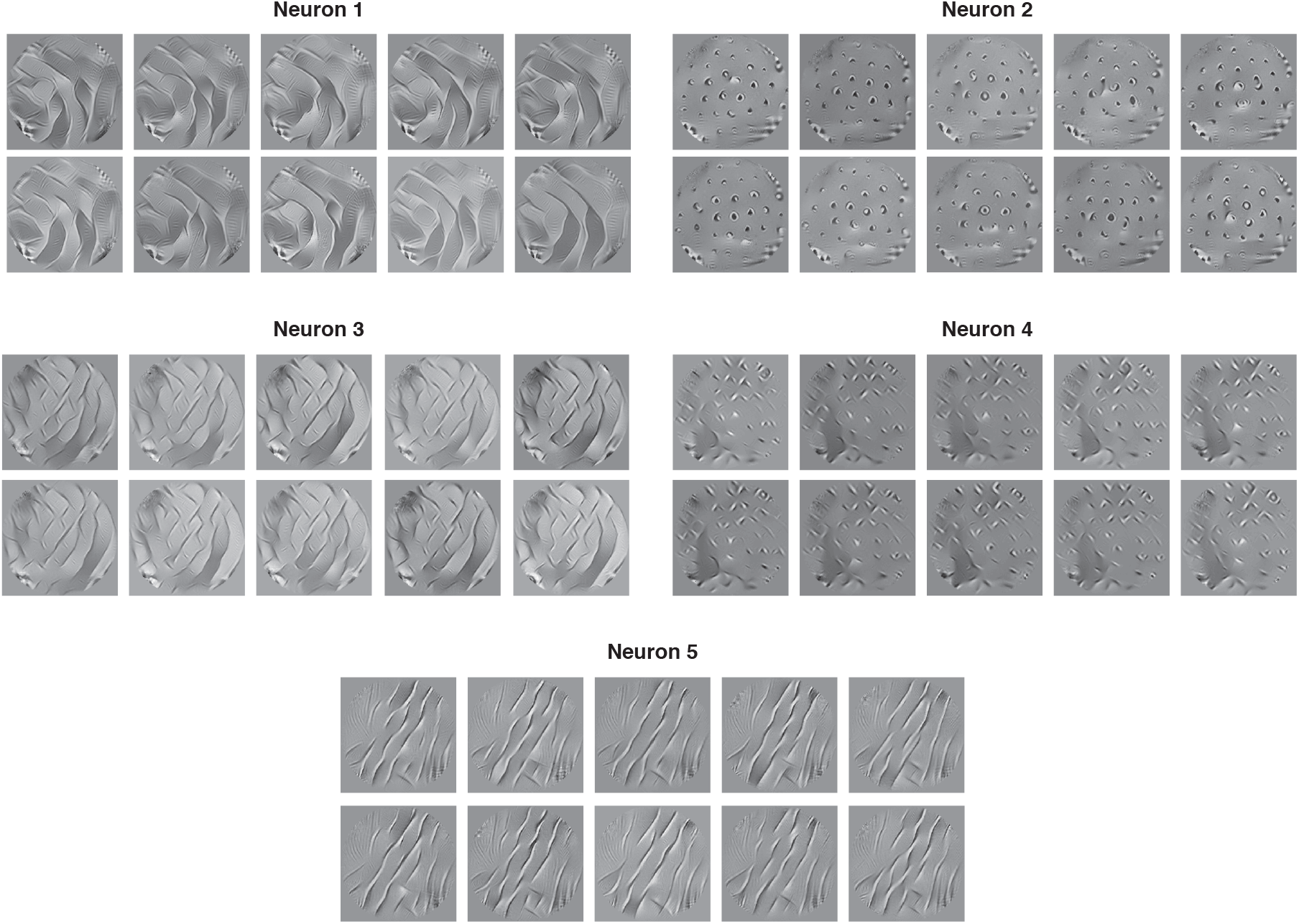
Smooth DeepTune images with 10 different random initializations for five neurons.

#### Stability across 18 models

The DeepTune images from all of the 18 models studies in this paper has stable patterns for each neuron. To construct the 18 models, we have used 3 pre-trained convolutional neural networks (AlexNet, VGG, and GoogleNet). From each network, we use either two, three, or four layers to extract features from images in neuroscience experiments. These features predict the spike rates of each neurons using a regularized linear regression. We use both *l*_2_ (ridge regression) and *l*_1_ (LASSO) regularizations. This results in 18 models for each neuron (3 networks, 3 layers, and 2 regression model). Figure 15 shows DeepTune images from each of these 18 models for two neurons. The stable pattern among these 18 DeepTunes should be interpreted as the pattern that activates the neuron.

**Figure 15:**
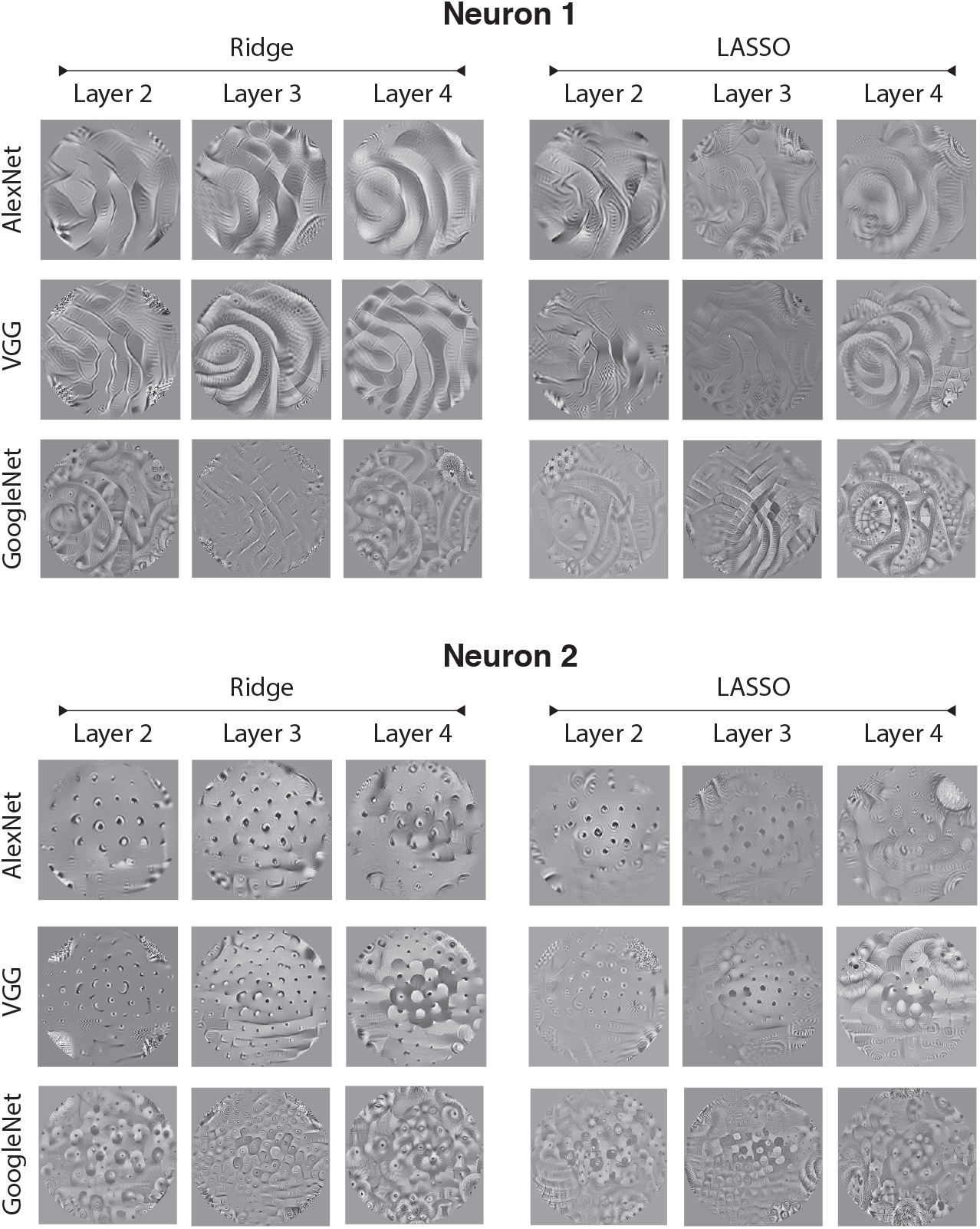
Stability of the interpretable patterns in DeepTune images for neurons 1 and 3 across 18 models. Three of the DeepTune processes for LASSO did not converge.

#### Identification matrix across models 1, 2, and 3

To demonstrate this diversity, we first compute DeepTune images for each neuron and then construct a response identification matrix. For each DeepTune image image, we compute the responses from each of the 71 neuron models to it and plot them together in the identification matrix in the top heatmap plot in Figure 16. The DeepTune image for each neuron has the highest response to model of that neuron compared to other neurons in the population (diagonal line visible in Figure 16). No pairs of columns looks exactly identical which is an evidence that the 71 neurons’ response properties are diverse. We also study the stability of this observation by feeding Deeptune images generated from VGG and GoogleNet models to AlexNet-based model. Figure 16 the middle and bottom heatmap plots illustrates the responses of neuron models from AlexNet layer 2 to DeepTune images generated by VGG and GoogleNet layer 2 models. Ridge regression have been used in all of the models. The heatmaps in Figure 16 contain clear diagonal patterns, showing the DeepTune images are stable across models. This observation, quantitatively confirms the visually observed stability of DeepTune images.

**Figure 16:**
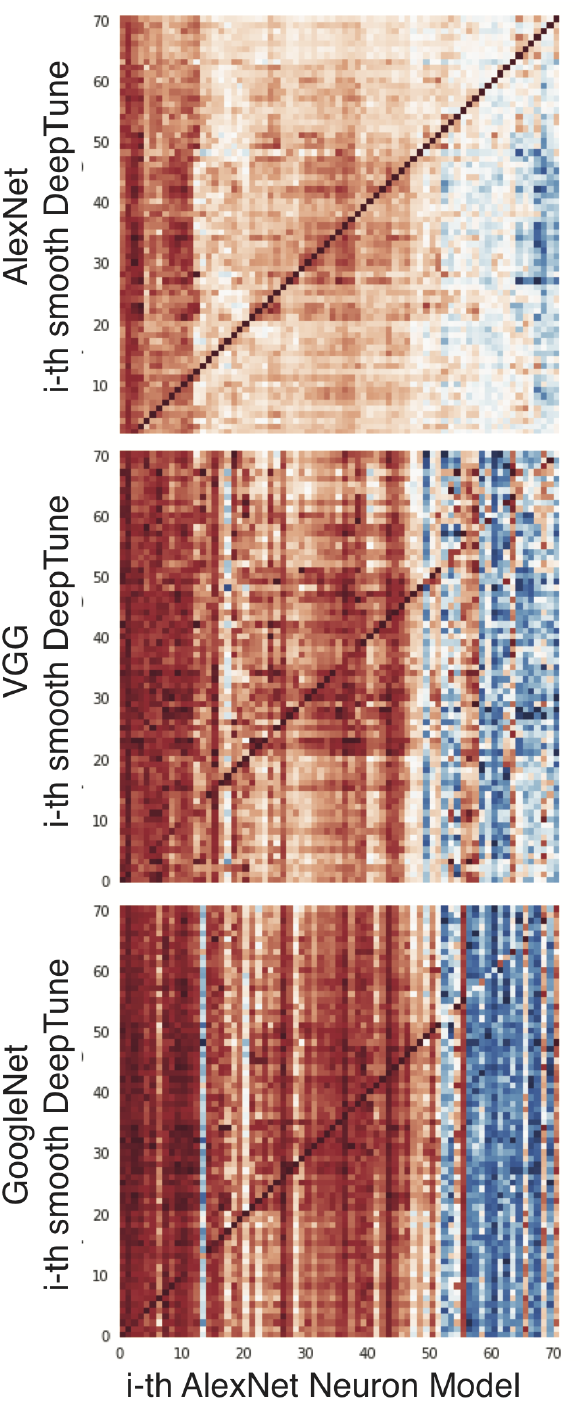
Smooth DeepTune images’ responses across models. Smooth DeepTune images from layer 2 of AlexNet, VGG and GoogleNet are generated for each neuron. These images are fed into our prediction model based layer 2 feature of AlexNet. All three heatmaps contain clear diagonal pattern, showing the DeepTune images are stable across models. This observation, quantitatively confirms the visually observed stability of DeepTune images.

#### Stability of Selected Features and Weight-Maps

In this section, we investigate the stability of CNN features selected by each neuron across different models. First, we visualize the top selected features and show that these features have stable visualization across models. Then, we use the regression coefficients in models to identify the model-inspired receptive field for each neuron. This is achieved by visualizing heatmaps of average regression coefficients across all features corresponding to each location in image.

#### Stability of top selected features across four main models for four neurons

Our model for each neuron consists of a CNN-based feature selection module and a linear regression model to predict the neuron spike rate from those features. Figure 17 shows that these features have stable visualization across models. For neurons 2, 3, 4, and 5, we visualize the filters representing top two selected features. Each box with 9 image patches visualizes a filter in the CNN. To visualize the filter, we feed a million random natural images (from AlexNet dataset) to the network and show the top 9 image patches that activate the filter. For each model the left box corresponds to the top filter and the right box corresponds to the second top filter selected by neuron. The patterns are stable across all four models. For neuron 2, both top and second top filters for four models are selective to blob-like patterns. CNN filters selected by Neuron 3 like both curvatures and diagonal edges with 45 deg patterns. Neuron 3 prefers filters selective to corners and edges in both diagonals, however, no curvature filter is selected by this neuron. Neuron 5 is consistently selecting filters responsive to diagonal patterns in 45 deg. Similar observation holds for other V4 neurons on the population.

**Figure 17:**
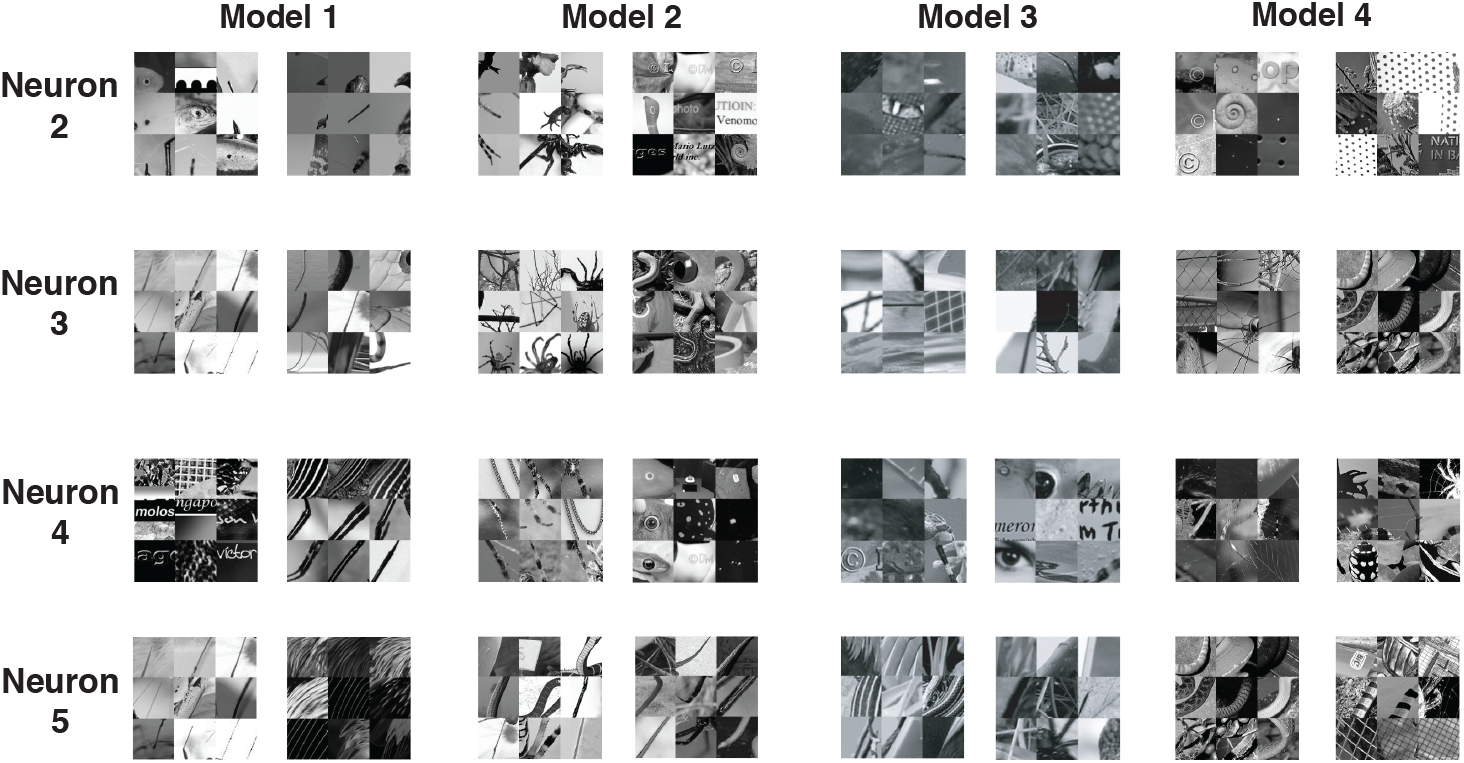
Stability of top selected CNN features selected by each neuron across four main models. Each box visualizes a filter representing the feature in the CNN. To visualize the filter, we feed a million random natural images (from AlexNet dataset) to the network and show the top 9 image patches that activate the filter.

#### Stability of weight-maps across four main models

In this section, we study the stability of model-inspired spatial receptive fields for each neuron. The CNN features extracted from images have spatial structure due to nature of convolution operation. That is, each feature corresponds to a location on the image. Consequently, the regression coefficients mapping these features to neuron spike rates have similar spatial structure. After fitting the predictive models for each neuron, we estimate a model-inspired receptive field for each neuron, by averaging the regression coefficients in each location across different filters. The heatmap of average regression coefficients for each location on the image represents the importance of that location for the neuron. Figure 18 shows these weight-maps for four neurons and four models. The weight-maps are stable across models. For neuron 2, the features in the center leaning to right side of the image are selected by the neuron. Neuron 3 and 5 are selective to features in a diagonal location. Neuron 4 prefers features in a cross-like location.

**Figure 18:**
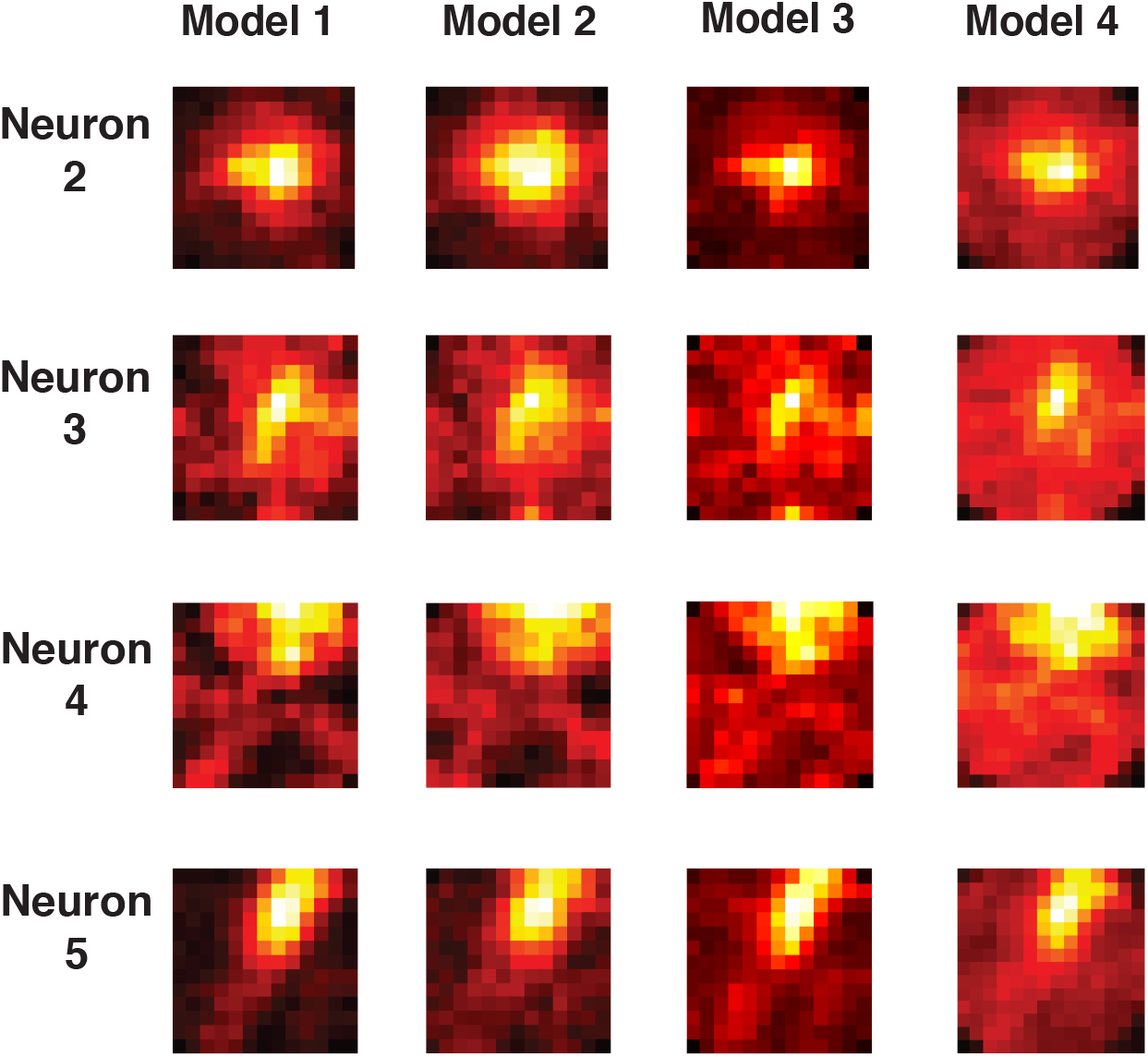
Stability of average model weight-maps across four main models. These weight-maps estimates the model-inspired receptive filed. After fitting the predictive models for each neuron, we estimate a model-inspired receptive field for each neuron, by averaging the regression coefficients in each location across different filters. Each row corresponds to one neuron. Each column corresponds to a model. The weight-maps are stable across models for each neuron.

**Figure 19:**
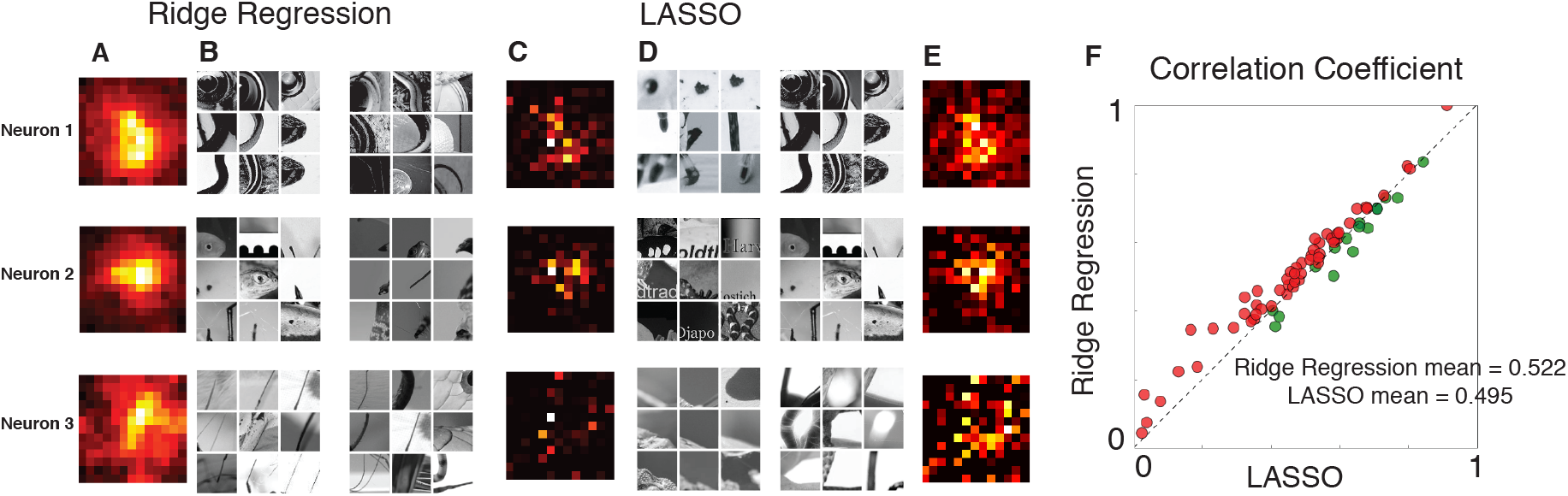
Comparison of Lasso and Ridge feature selection. Ridge and Lasso give similar prediction performance. Lasso in general selects a smaller number of features (751 features in average) included in the set of features that Ridge selected (total 377,000 features). The top CNN filters that Lasso selected are similar to that of the Ridge regression.

#### Stability LASSO vs Ridge

Inspecting raw coefficients from models learned by Lasso is problematic, however, due to the instability of the Lasso selected features. Particularly in cases where regressors are highly correlated (as is the case with features extracted from a CNNs), the model selection performed by Lasso may be inconsistent [52]. To overcome this issue and focus on the truly salient features for a particular neuron, we performed a stability analysis using 10-fold cross validation: the model was refit on each of the 10 perturbed datasets, and then the sets of selected variables were intersected. Model coefficients on this set were then averaged and used as a basis for our analysis. This is similar to the method introduced in [3], except we use cross validation instead of bootstrap resampling.

### D Population Analysis of V4 Neurons

In this section, the preferred and non-preferred DeepTune images are visualized for all of the 71 V4 neurons in the population.

#### Preferred DeepTune images for all 71 V4 neurons

Figure 20 shows the preferred DeepTune images for all of the 71 neurons under study in visual area V4. The model used here is AlexNet layer 2 with ridge regression. Refer to the main text for a discussion on the diversity of the patterns preferred by V4 neurons.

**Figure 20:**
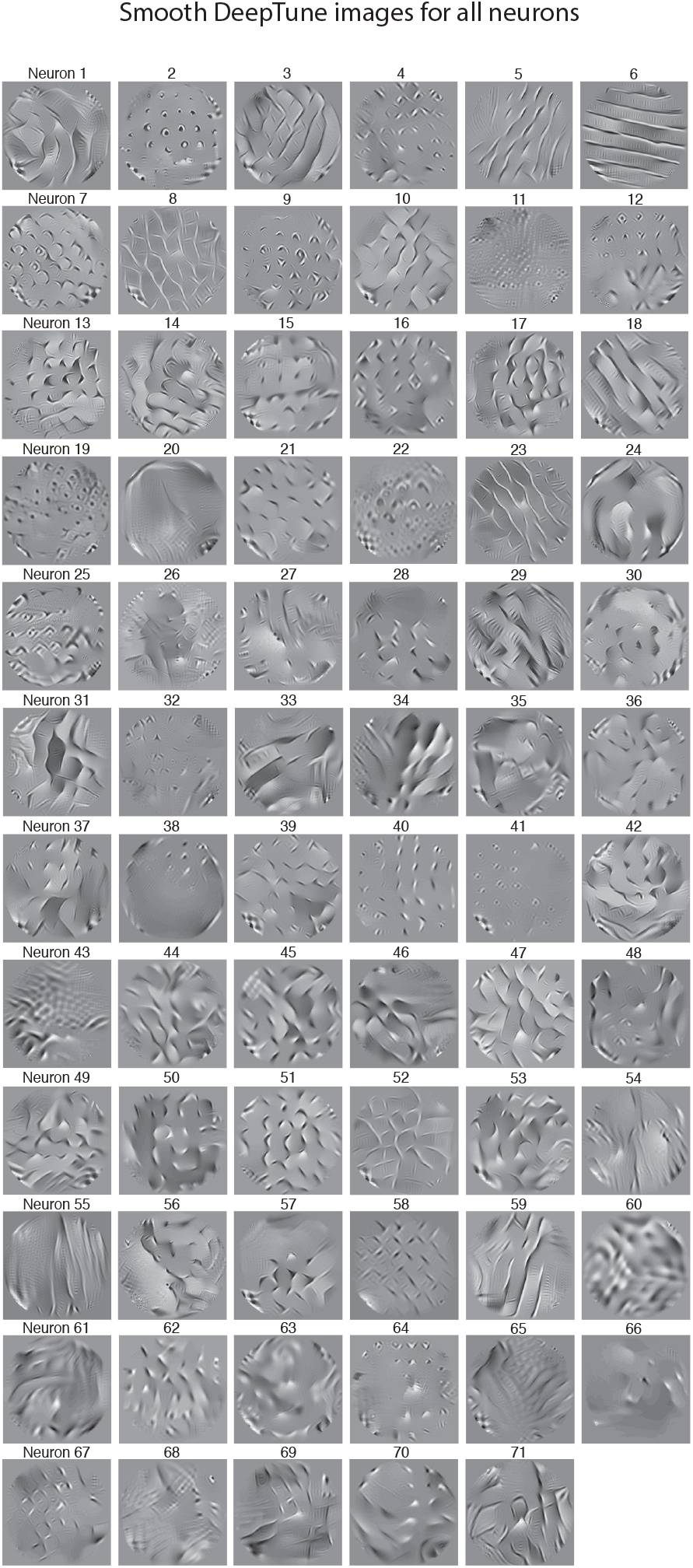
Smooth DeepTune images for all 71 V4 neurons, based on AlexNet

#### Non-preferred DeepTune images for all 71 V4 neurons

Figure 21 illustrates the non-preferred DeepTune images for all of the 71 neurons. The model used here is AlexNet layer 2 with ridge regression. Most of the neurons have weak patterns in their non-preferred DeepTune image. For some of the neurons, the pattern is stronger. In the main text of chapter 3, we present a detailed discussion on the interpretation of patterns in non-preferred DeepTune images.

**Figure 21:**
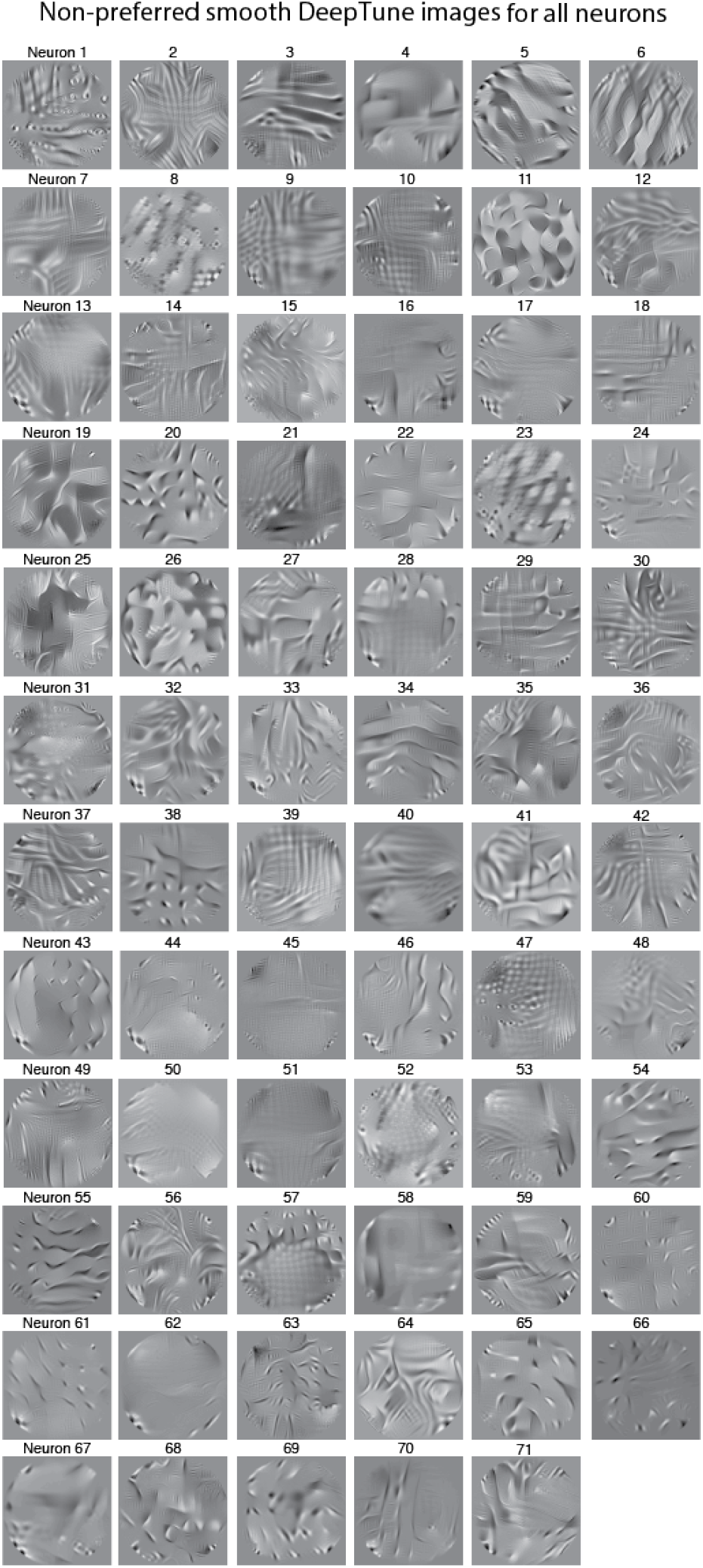
Inhibitory DeepTune images for all 71 V4 neurons, based on AlexNet

**Figure 22:**
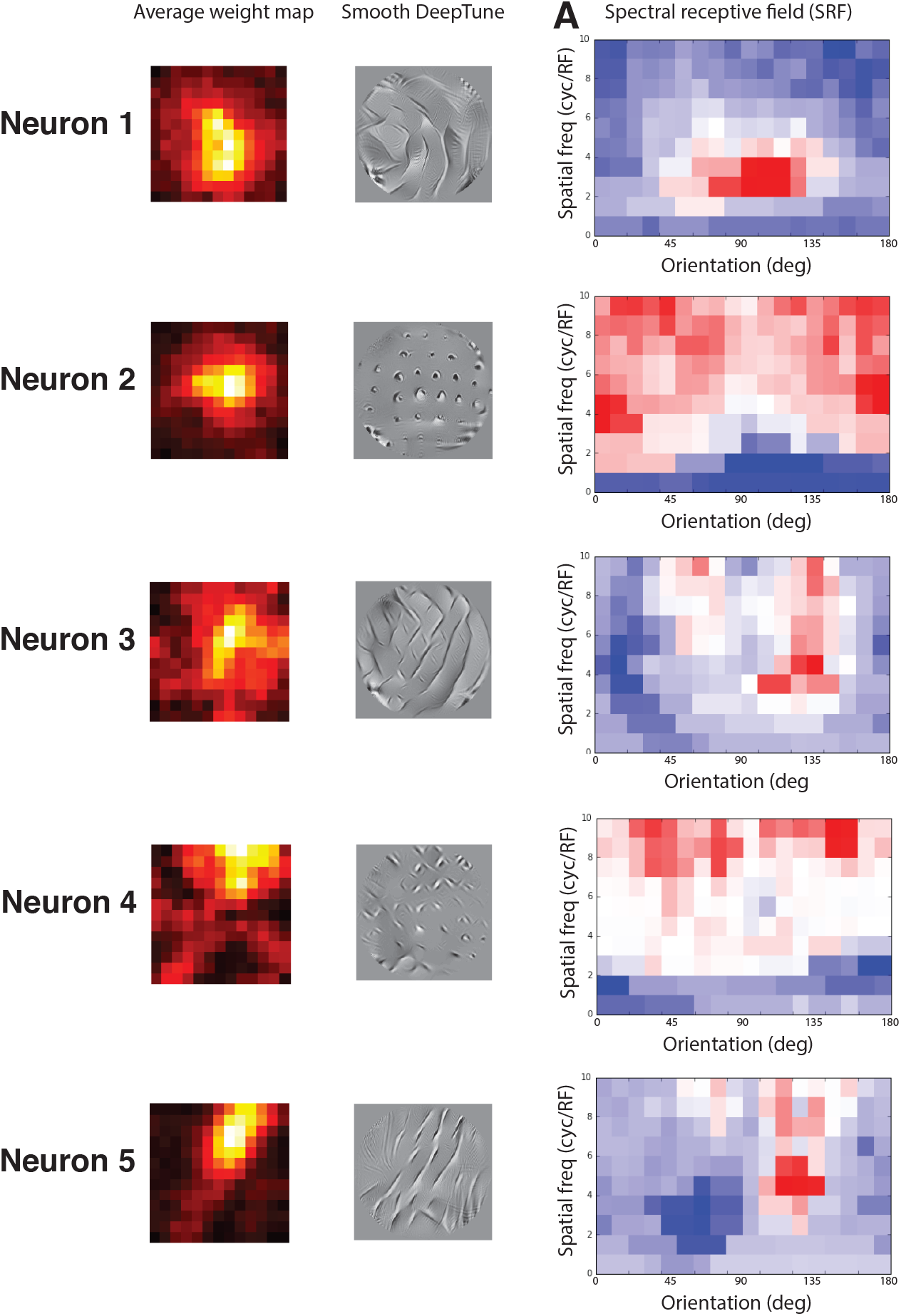
Consistency of the average weight map and DeepTune images with spectral receptive field (SRF) [8].

### E Analysis of Our Data based on Previous Methods

#### Spectral Receptive Field (SRF)

It has been shown in [8] many V4 neurons have more than one excitatory orientation tuning peak. Bimodal orientation tuning explains previous observations of selectivity for sharp corners [27]. Curvature or corner patterns will result in Bimodal orientation tuning in V4. We show via DeepTune that a large part of V4 neurons share this property and the result is consistent with that obtained by the spectral receptive field (SRF) [8].

### F Principal Component Analysis

Each V4 neuron model corresponds to a point in *p*-dimensional coefficient space; we can investigate the population of V4 neurons by examining their relative positions in this space. However, because *p* is very large (in the case of models based on N2 of AlexNet, *p* = 389, 376) direct analysis of the coefficient vectors is impossible due to the curse of dimensionality. First, we perform *ℓ*^2^ pooling of coefficient values across space and time delays to yield a single impact value for each of the filters in layer N2 of AlexNet. This gives a 256-dimensional representation, where each dimension corresponds to a single filter. Next, we perform principal components analysis (PCA) of the 71 points (each corresponding to a single V4 neuron) in this 256-dimensional space. PCA finds a set of linear transformations that capture a large proportion of the variance of the vectors. An examination of the coefficients of the loading vectors reveals that the first several principal components delineate several recognizable image features. The first principal component specifies whether neuron is selective to horizontal and vertical patterns. The second principal component delineates low-frequency patterns vs. dense blobs. The third principal component delineates diagonal vs non-diagonal features.

Figure 23.A shows the plot of the 71 V4 neurons according to their values in the first two principal components. Each neuron is shown via its DeepTune image. The color of DeepTune image borders is proportional to the third principal component with red being the highest PC value and blue the lowest. The neurons with highest values in PC 1 are selective to Figure 23.B illustrates the 71 V4 neurons according to other principal components. The coefficients of the loading vectors for first three principal components are shown in Figure 23.C. For the top coefficients, the corresponding filter is visualized by top 6 image patch that activate that filter. These image patches are found by feeding one million random image to the CNN and selecting the patches with highest filter response.

**Figure 23:**
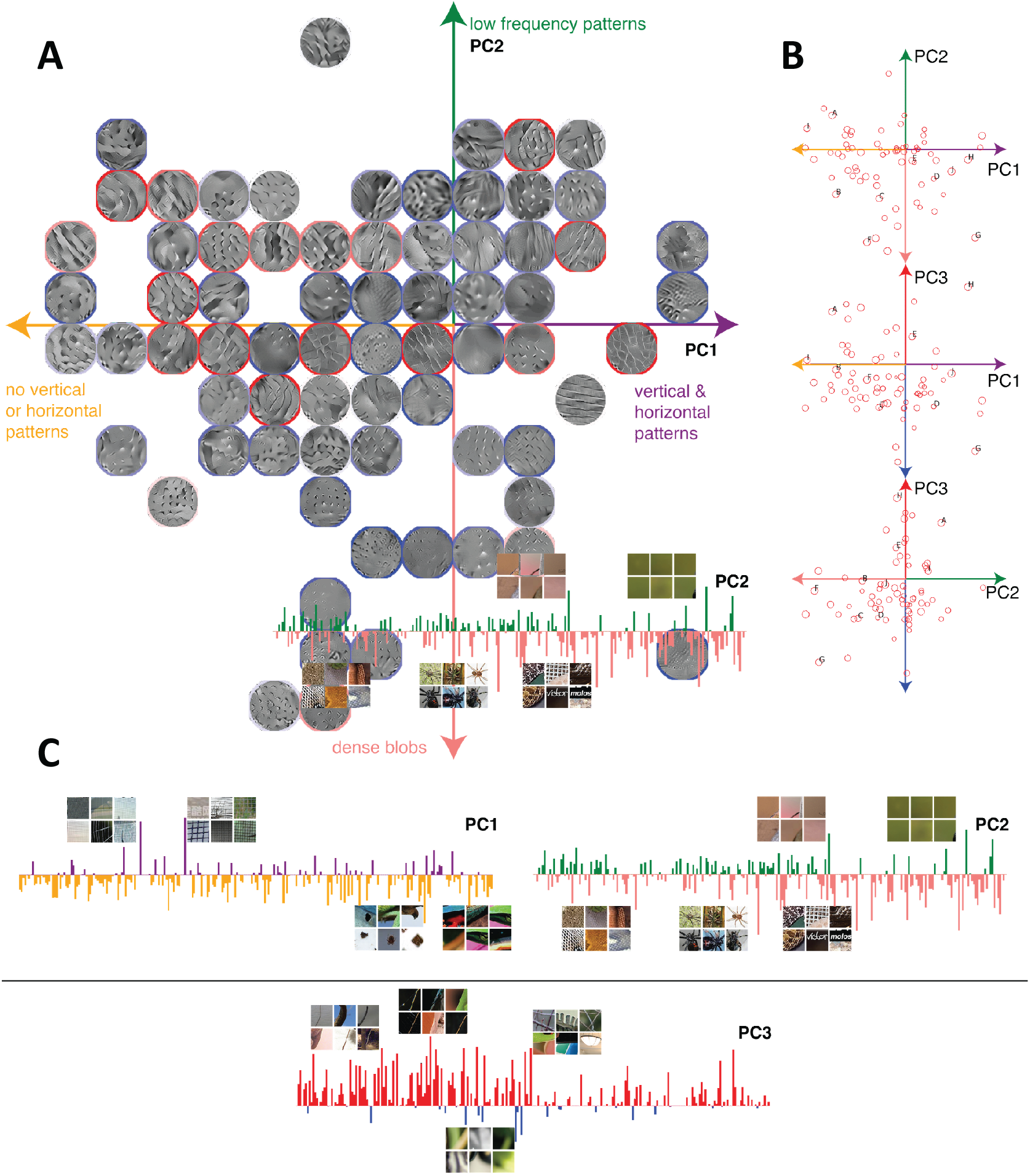
Principle components analysis of V4 neuron’s population. **A.** 71 V4 neurons according to their values in the first two principal components. To compute the principal components, we perform *ℓ*^2^ pooling of coefficient values across space and time delays to yield a single impact value for each of the filters in layer N2 of AlexNet. This gives a 256-dimensional representation, where each dimension corresponds to a single filter. Then, we perform principal components analysis (PCA) of the 71 points (each corresponding to a single V4 neuron) in this 256-dimensional space. Each neuron is shown via its DeepTune image. The color of DeepTune image borders is proportional to the third principle component with red being the highest PC value and blue the lowest. **B.** 71 V4 neurons according to other pairs of principal components. **C.** Coefficients of the loading vectors for the top three principal components. For the coefficients with highest values, the corresponding filter is visualized by top 6 image patch that activate that filter. These image patches are found by feeding one million random image to the CNN and selecting the patches with highest filter response.

### G Additional Figures

#### Responses of our model to hand-crafted stimuli

In this section, we investigate the response of our CNN-based neuron models to polar, hyperbolic, and Cartesian gratings. We manually create images in each category based on the equations give in [10]. The response of the V4 models based on second layer of AlexNet with Ridge regression are computed for each of these hand crafted images. Figures 24, 25, and 26 show the responses of three neurons to these images. The smooth DeepTune image is also shown in each figure. The hand crafted images selected by the model are consistent with the patterns visible in the DeepTune image.

**Figure 24:**
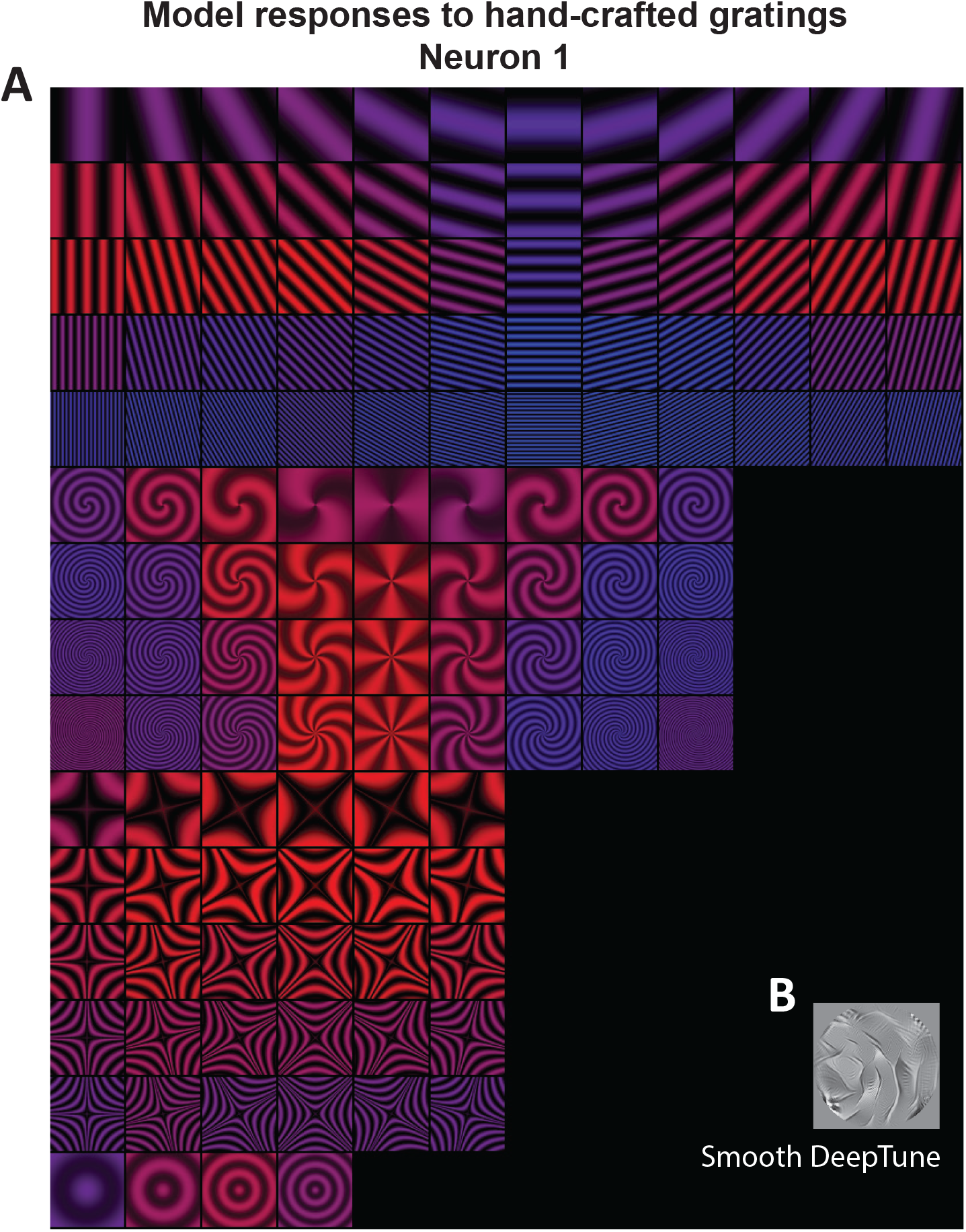
A. Responses of neuron 1 model to polar, hyperbolic, and Cartesian gratings. The gratings in red and blue correspond to excitatory and inhibitory stimulus, respectively. B. Smooth DeepTune of neuron 1

**Figure 25:**
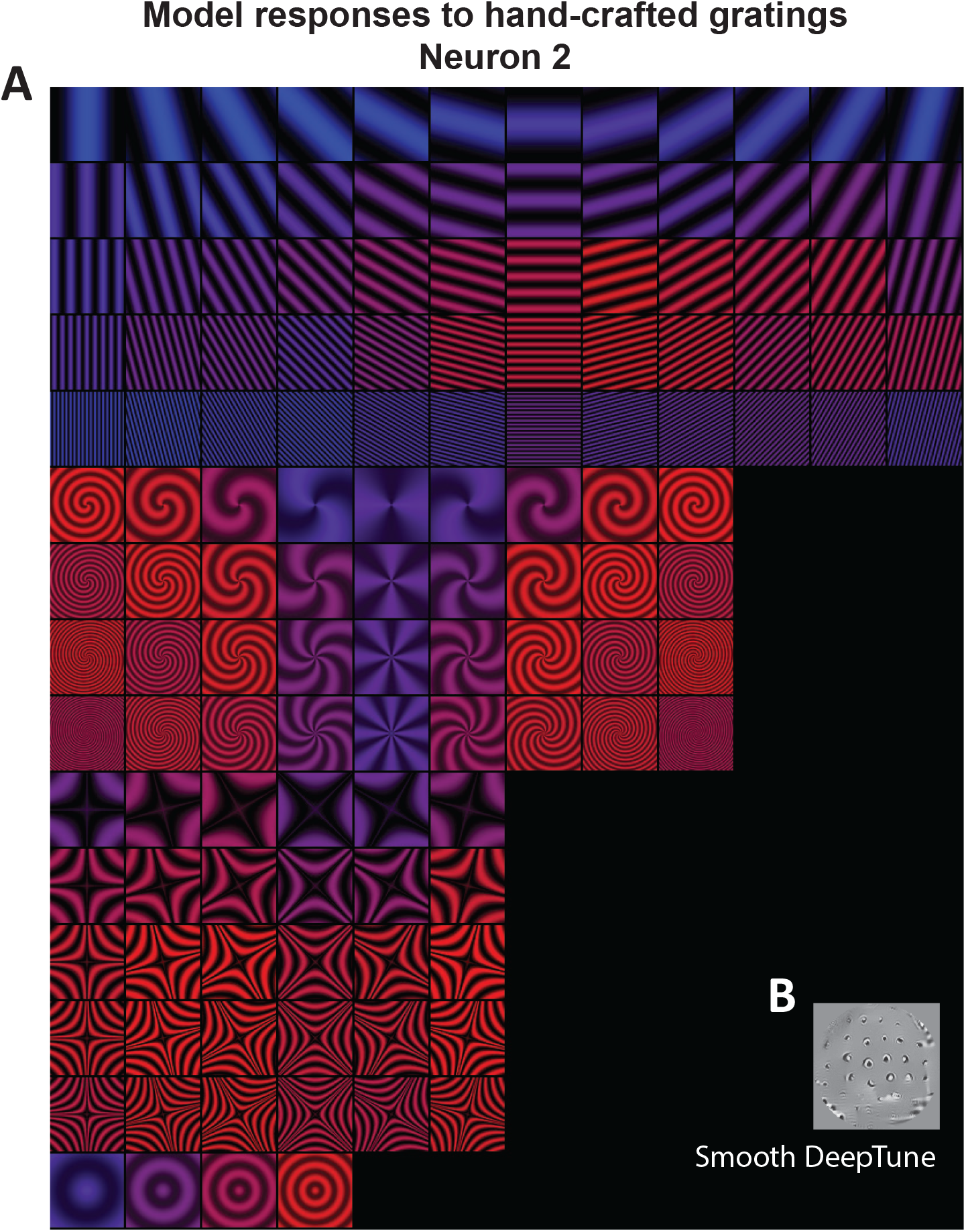
A. Responses of neuron 2 model to polar, hyperbolic, and Cartesian gratings. The gratings in red and blue correspond to excitatory and inhibitory stimulus, respectively. B. Smooth DeepTune of neuron 2

**Figure 26:**
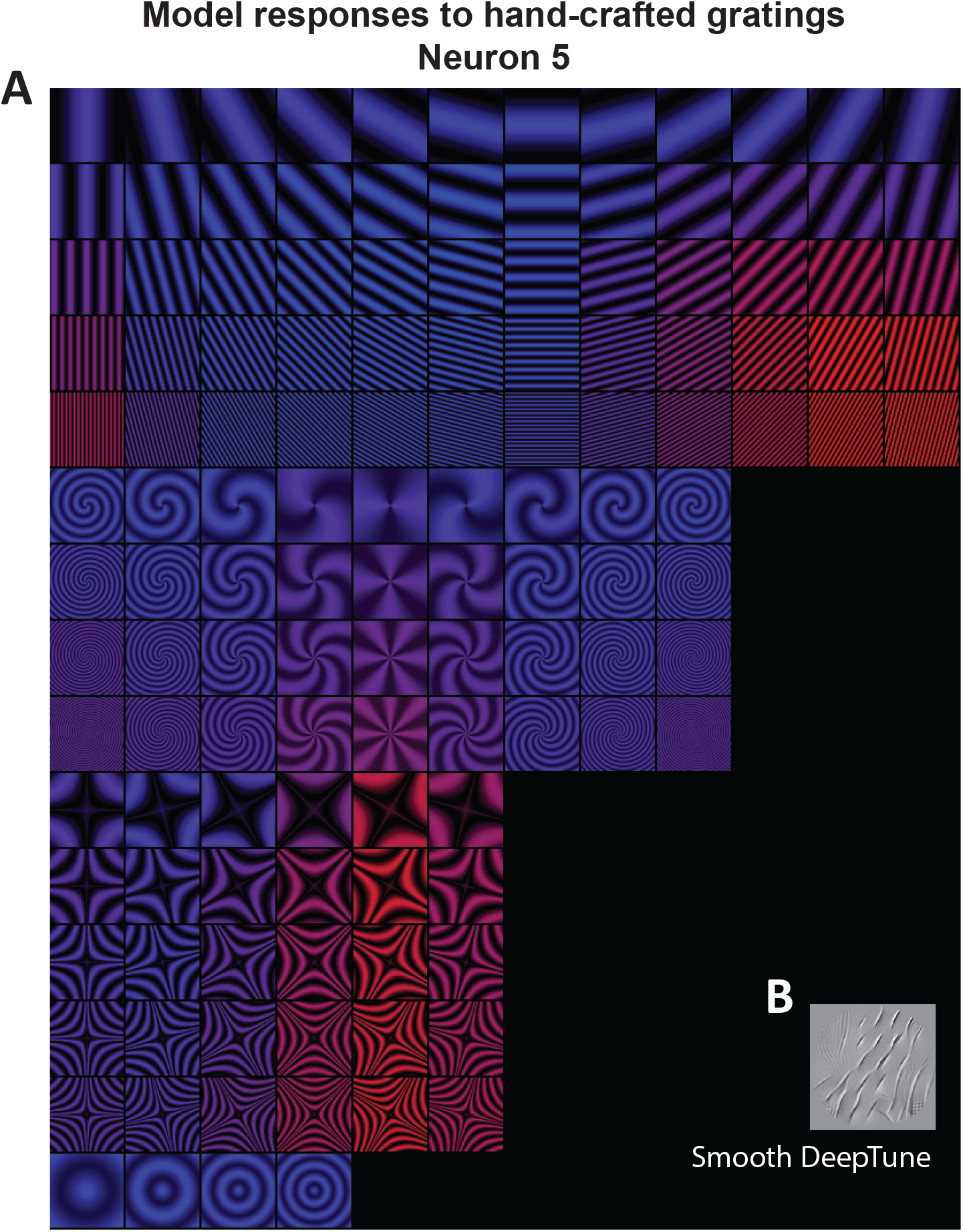
A. Responses of neuron 5 model to polar, hyperbolic, and Cartesian gratings. The gratings in red and blue correspond to excitatory and inhibitory stimulus, respectively. B. Smooth DeepTune of neuron 5

**Figure 27:**
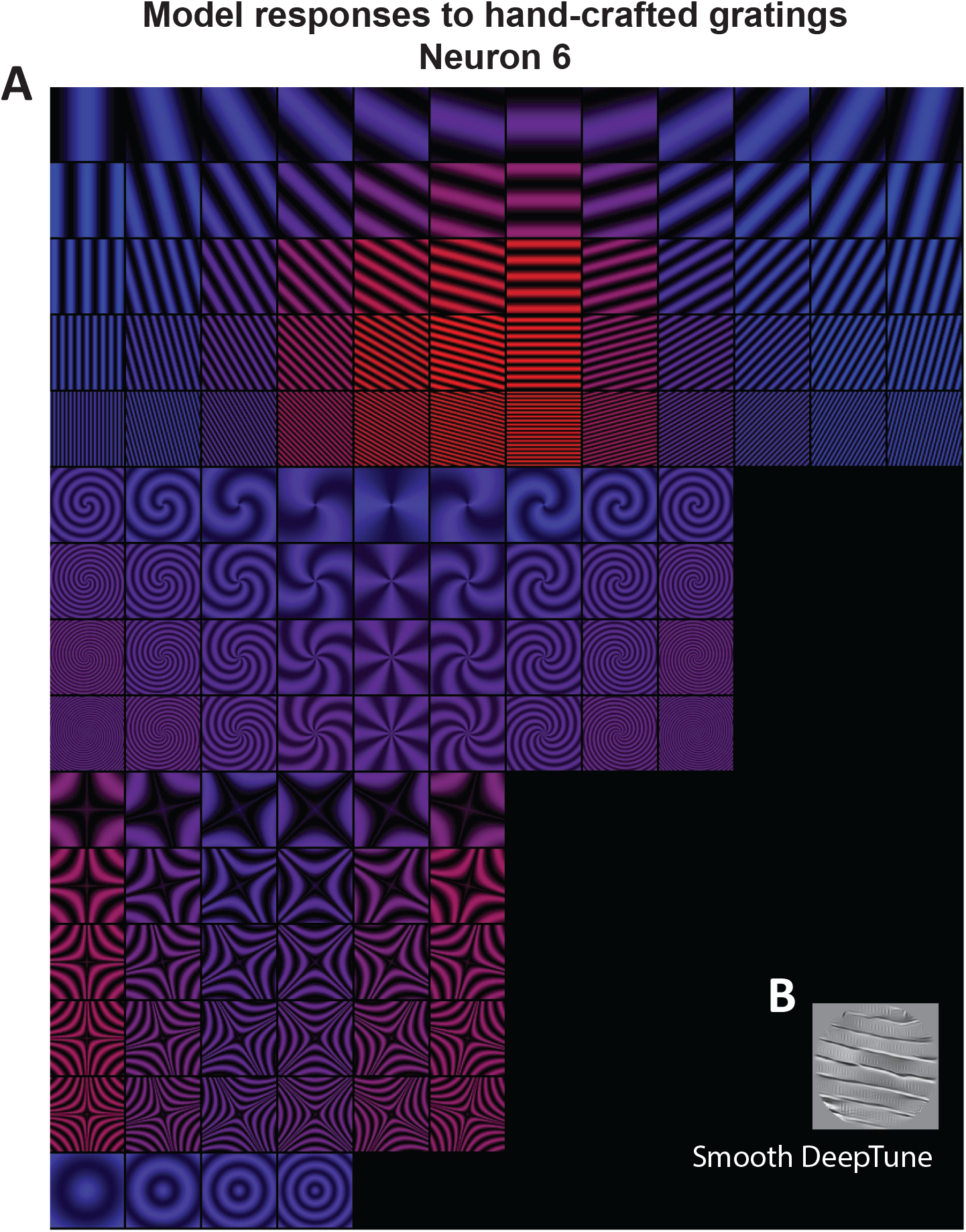
A. Responses of neuron 6 model to polar, hyperbolic, and Cartesian gratings. The gratings in red and blue correspond to excitatory and inhibitory stimulus, respectively. B. Smooth DeepTune of neuron 6

